# Competition for light can drive adverse species-composition shifts in the Amazon Forest under elevated CO_2_

**DOI:** 10.1101/2023.07.03.547575

**Authors:** Jaideep Joshi, Florian Hofhansl, Shipra Singh, Benjamin D. Stocker, Åke Brännström, Oskar Franklin, Carolina C. Blanco, Izabela F. Aleixo, David Lapola, Iain Colin Prentice, Ulf Dieckmann

## Abstract

The resilience of biodiverse forests to climate change depends on an interplay of adaptive processes operating at multiple temporal and organizational scales. These include short-term acclimation of physiological processes like photosynthesis and respiration, mid-term changes in forest structure due to competition, and long-term changes in community composition arising from competitive exclusion and genetic trait evolution. To investigate the roles of diversity and adaptation for forest resilience, we present Plant-FATE, a parsimonious eco-evolutionary vegetation model. Tested with data from a hyperdiverse Amazonian terra-firme forest, our model accurately predicts multiple emergent ecosystem properties characterizing forest structure and function. Under elevated CO_2_ conditions, we predict an increase in productivity, leaf area, and aboveground biomass, with the magnitude of this increase declining in nutrient-deprived soils if trees allocate more carbon to the rhizosphere to overcome nutrient limitation. Furthermore, increased aboveground productivity leads to greater competition for light and drives a shift in community composition towards fast-growing but short-lived species characterized by lower wood densities. Such a transition reduces the carbon residence time of woody biomass, dampening carbon-sink strength and potentially rendering the Amazon Forest more vulnerable to future climatic extreme events.

## 1 Introduction

Intact tropical forests currently act as global carbon (C) sinks, partially offsetting the effects of deforestation and fossil fuel burning^1, 2^. However, since the 1990s, despite increasing atmospheric CO_2_ concentrations, the tropical forest C sink has saturated in Africa and decreased in Amazonia^3^. While CO_2_ fertilization may potentially increase the productivity of terrestrial ecosystems, it has been argued that these gains could be diminished by higher tree mortality rates arising from higher temperatures^4^, growth-lifespan tradeoffs^5, 6^, and recurrent climatic extreme events such as droughts^7–9^. Nutrient availability can additionally constrain the effects of CO_2_ fertilization in the short term, according to ecosystem manipulation experiments^10^ and vegetation models^11^. However, it has been hypothesized that plants can overcome low nutrient availability in the long run by investing more resources in nutrient acquisition (e.g., via greater C allocation to fine roots^11, 12^) or by utilizing nutrients deposited from anthropogenic sources^13^. Furthermore, changes in CO_2_ or nutrient concentrations can alter species compositions or drive adaptive changes in species traits, which may fundamentally affect vegetation carbon stocks and community functioning^14–16^. For example, several studies have reported a consistent decline in tree wood density over the past century^17–19^, dampening the increase in plant biomass that could have resulted from increased volumetric growth. Therefore, predictions of ecosystem responses to climate change on multi-decadal to centennial timescales must also account for simultaneous changes in tree community composition. However, standard DGVMs, which group vegetation into a fixed set of ‘plant functional types’ (PFTs), lack sufficient representation of functional diversity and demographic dynamics to predict such changes.

For accurately predicting ecosystem responses to elevated CO_2_ and climate change, we need models that can account for adaptations encompassing multiple temporal and organizational scales. These are (1) short-term acclimation of plastic physiological traits in response to environmental change, (2) mid-term demographic changes in vegetation structure and species abundances, and (3) long-term evolution of the community structure via (a) competitive exclusion of less fit species and (b) gradual evolution of species’ genetic traits by natural selection. While current ‘dynamic global vegetation models’ (DGVMs) fall short of one or more of these requirements, emerging next-generation vegetation models^20, 21^ are beginning to address some of these shortcomings, as described in further detail below.

First, many current DGVMs typically use empirically derived response functions for modelling short-term physiological responses. This approach does not account for trade-offs faced by plants and thus does not scale well to out-of-sample environmental conditions. This causes their predictions to be particularly sensitive to model assumptions and uncertain parameter values^11^, resulting in large uncertainties and poor agreement among models^22, 23^. In recent years, eco-evolutionary optimality (EEO)^24, 25^ has emerged as a powerful principle for modelling physiological acclimation. The EEO approach naturally accounts for trade-offs, allowing for simpler and more robust representations of physiological processes. For instance, optimality-based models, using fewer parameters than their empirical counterparts, have successfully predicted the environmental dependencies of photosynthesis rates^26–28^, hydraulic traits^29, 30^, and leaf lifespans^31^.

Second, many current DGVMs lack representation of size structure, which is critical for capturing size-structured competition (e.g., for light) and size-dependent demographic trade-offs. Indeed, size-dependent strategies make plant size the second most important axis of variation in plants, after the leaf-economics axis^32^. A new class of models, called vegetation demographic models (VDMs), is beginning to incorporate size-structure^21^. Such models predict qualitatively different community-level properties compared to similar models without size structure^33^. Furthermore, incorporating size structure allows for modelling the continuous acclimation of physiology to changing biotic environments that trees experience as they grow from the understorey into the canopy. Such translation of short-term acclimating responses to mid-term demographic rates provides a basis for predicting carbon stocks, demographics, succession^34^, diversity, and coexistence^35^.

Third, most DGVMs, even those that incorporate size-structured demographics, rely on PFTs to represent different vegetation types. This limits their capacity to account for functional diversity^36–40^, a crucial ingredient for the resilience of hyperdiverse ecosystems. To that end, DGVMs are increasingly adopting a trait-based approach^20, 41, 42^, where species are represented as points in a multidimensional trait space. Since traits are subject to natural selection, such an approach is also appropriate for predicting the long-term evolution of species. At the community level, competition between individuals and feedbacks between the population structure and the environment may lead to non-optimal outcomes^43^, rendering the EEO approach ineffective. In such cases, game-theoretic approaches or evolutionary dynamics theory can be used to predict evolutionarily stable community-level properties. For example, models utilizing this approach have succeeded, albeit in limited settings, in predicting evolutionarily stable strategies for biomass allocation^44^, crown-shapes^45^, timings of budburst^46^, and community assembly^35^. Currently, only a few models (e.g., ref.^47^) incorporate trait evolution, typically using an agent-based approach.

Here, we unify four key emerging concepts and tools into a parsimonious eco-evolutionary vegetation model – Plant-FATE (Plant Functional Acclimation and Trait Evolution). These are: (i) EEO-based descriptions of physiological acclimation, (ii) a multidimensional trait-based representation of functional diversity, (iii) size-structured competition for light and vegetation demographics, and (iv) evolution of species traits by natural selection. We calibrate the model for a mature central-eastern Amazonian terra-firme forest. We then use the model to elucidate the responses of such a forest to elevated atmospheric CO_2_ concentrations. Specifically, we ask three questions: (1) What are the processes that affect community-level responses to elevated CO_2_, and what are the timescales of these responses? (2) How does size-structured competition for resources (such as light) affect community-level responses? (3) How do changes in belowground resource availability, and resulting physiological acclimation, affect community-level responses?

## 2 Plant-FATE model summary

Plant-FATE is a trait-size-structured eco-evolutionary vegetation model that leverages the principles of natural selection to predict multidimensional and multiscale adaptive responses of forest ecosystems to climate change. Trait structure allows for representing functional diversity and capturing trait-mediated trade-offs in plant function that ultimately determine their fitness. Size structure allows for modelling size-dependent demographics and competition for light. To model the short-term acclimation of physiological processes such as CO_2_ assimilation, respiration, and turnover, we employ eco-evolutionary optimality principles. To model mid-term (and long-term) changes in species abundances resulting from vegetation demographics and competition for light, we use a physiologically structured population modelling (PSPM) framework based on partial differential equations. To model the long-term evolution of species’ genetic traits, we use evolutionary dynamics theory. Forced with four environmental parameters (temperature, light intensity, vapour pressure deficit, and soil moisture), Plant-FATE predicts the fate of multi-species communities. It predicts the temporal dynamics of the fluxes of CO_2_ and water, forest structure, and trait distributions. Below, we describe key model features that are novel compared to existing models. Equations describing the main processes can be found in the methods section, and a full description and derivation of all model components can be found in the Supplementary Information.

### Plant physiology and its acclimation

To the extent that the latest theoretical developments allow, we use trait-based and EEO-based representations of three core processes (photosynthesis, leaf economics, and xylem construction), as described below.

1. Optimal photosynthesis. The acclimation of photosynthesis is the fundamental basis for predicting ecosystem productivity and adaptations at higher scales. Therefore, an accurate representation of the environmental dependencies of leaf-level photosynthesis is critical for reducing uncertainties downstream. To that end, we use the P-hydro model^28^, which predicts the acclimation of leaf photosynthetic capacities (*V*_cmax_ and *J*_max_), stomatal conductance (*g*_c_), soil-to-leaf water potential difference (Δ*ψ*), leaf-internal-to-external CO_2_ ratio (*χ*), and assimilation rate (*P*_c_). The strength of the P-hydro model is that it accounts for the biochemical and hydraulic controls of photosynthesis from first principles, and thus captures the observed environmental responses to CO_2_, vapour pressure deficit (VPD), light intensity, and soil moisture^28^.
2. Optimal leaf economics. In megadiverse forests such as the Amazon, shade tolerance is a crucial feature that maintains biodiversity by allowing short-statured species or juvenile trees to survive in the dark understorey. In our model, shade tolerance is not prescribed but emerges from an acclimation of leaf lifespan to the prevailing light environment. To model this acclimation, we use a modified version of the optimality-based theory of leaf economics developed in ref.^31^. The theory predicts that leaf lifespan is proportional to leaf mass per unit area (LMA) and inversely proportional to *V*_cmax_. We model a similar acclimation of fine-root lifespan based on observed effects of productivity and specific root length on fine-root longevity^48, 49^. As a result, evergreen trees in the understorey have lower leaf and fine-root turnover rates and, consequently, greater net production and higher chances of survival.
3. Optimal hydraulics. Plant water balance plays a critical role in regulating the rates of photosynthesis and drought-induced mortality. The hydraulic pathway consists of the xylem and outside-xylem segments, each with its own set of hydraulic traits. We utilize a simplified version of the theory of optimal xylem construction proposed in ref.^30^ to model the coordination of the xylem and outside-xylem segments, thus reducing the number of independent hydraulic traits.

### Vegetation demographics

Leaf-level photosynthesis is scaled up to the level of the whole tree by integrating leaf-level assimilation rate over the tree crown, accounting for the fact that different parts of the crown may experience different light levels. The assimilated carbon is allocated to various carbon pools (leaves, branches, trunk, coarse roots, and fine roots) using an extended version of the T-model^50^. The T-model is a model of biomass allocation in which all dimensional variables, such as height, crown area, stem mass, and root mass, scale with the stem diameter through scaling relationships. It needs fewer parameters than allometric models typically used in biomass allocation schemes. Moreover, each parameter is a measurable trait that encodes distinct life-history strategies. The T-model calculates biomass allocation to diameter growth and reproduction. Mortality rate is calculated from diameter, productivity, and wood density, using an empirically derived function. The emerging three demographic rates form the inputs for a physiologically structured population model that predicts the dynamics of the tree size distribution over time.

### Competition for light

Individual trees try to optimize their performance in the given local environment, but all trees in the population also collectively create or modify the local environment. Specifically, taller trees shade shorter trees, leading to competition for light. The vertical light profile is determined by the vertical distribution of leaves, which in turn depends on (a) the size distribution of trees in the population, (b) the vertical distribution of leaves within a tree crown, and (c) the spatial distribution of tree crowns. While the size distribution of trees is predicted by the demographic model, we use optimality arguments for the latter two distributions.

To model the distribution of leaves within the crown, we assume that branches are arranged to minimize self-overlap within the crown. However, each branch can have overlapping leaves, creating a potentially high within-crown leaf area index. To account for the spatial distribution of crowns in the forest canopy, we utilize an optimality idea called the perfect plasticity approximation (PPA)^51^.

The PPA states that trees position their crowns in canopy gaps to minimize overlap with other crowns by slightly flexing their stems. This overlap-minimizing behaviour of tree crowns leads to spontaneous emergence of discrete canopy layers, each of which contains crowns that experience similar light intensities and whose total crown area equals the ground area.

### Functional diversity

Plant-FATE uses a multidimensional representation of functional diversity. The community comprises multiple functional species, defined as unique combinations of genetic traits. To model the functional elasticity of species, we account for intraspecific variation in genetic traits and phenotypic plasticity in ontogenetic traits. Thus, species are represented by narrow Gaussian distributions in the multidimensional trait space, whose variance-covariance matrices capture intraspecific trait variation. In principle, all genetic (non-acclimating) traits should be used to characterize a species. However, in this work, we characterize species with only four traits, while all other traits are held constant across species (as specified in Table 1). These four traits are (i) LMA *λ*, (ii) wood density *ρ*_s_, (iii) maximum height *H*_m_, and (iv) xylem hydraulic vulnerability *ψ*_50x_. Although there is evidence of some levels of plasticity in these traits over decadal timescales, we treat them as genetic traits because of relatively high interspecific variance or their longer acclimation timeframes.

**Table 1.**
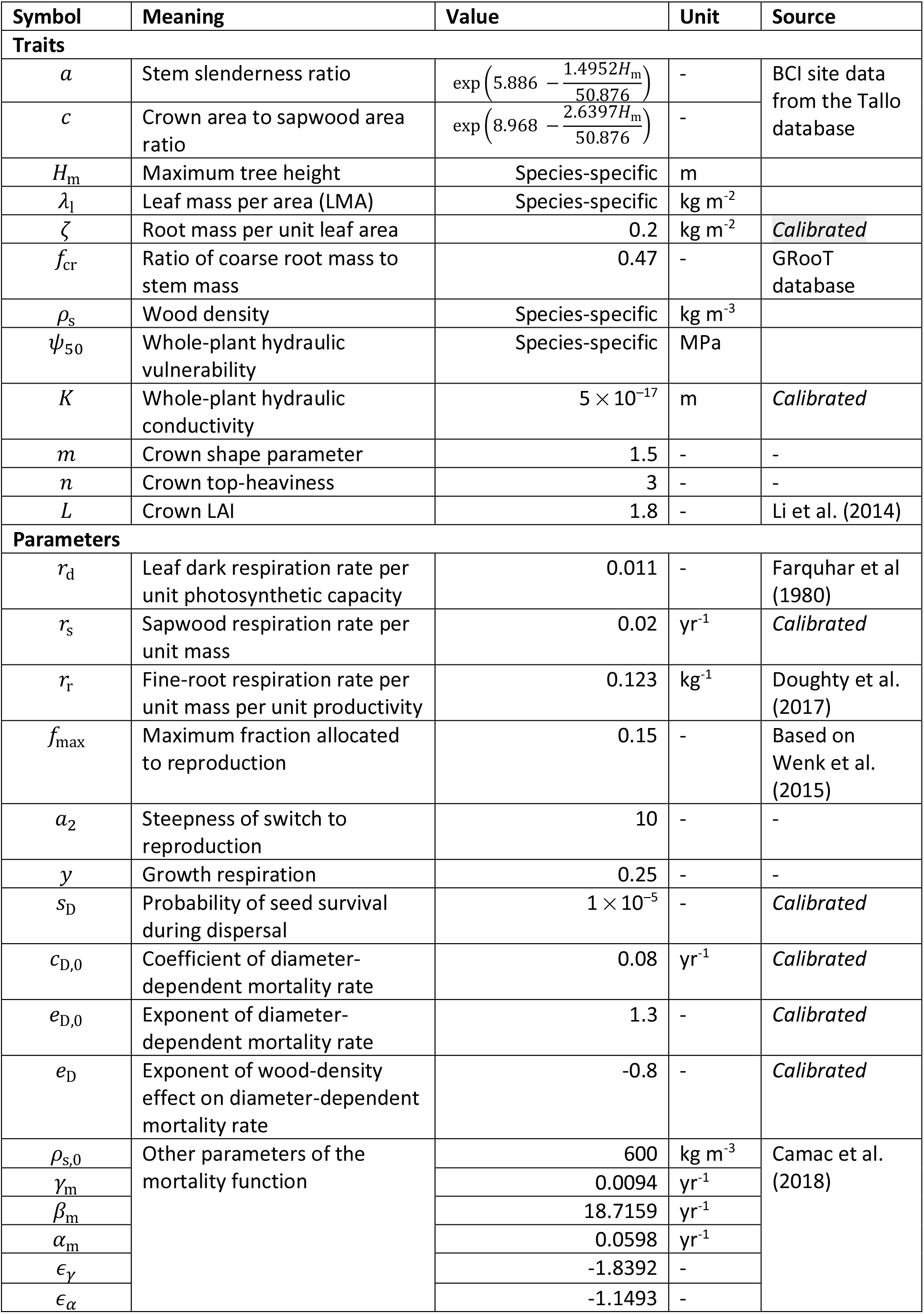
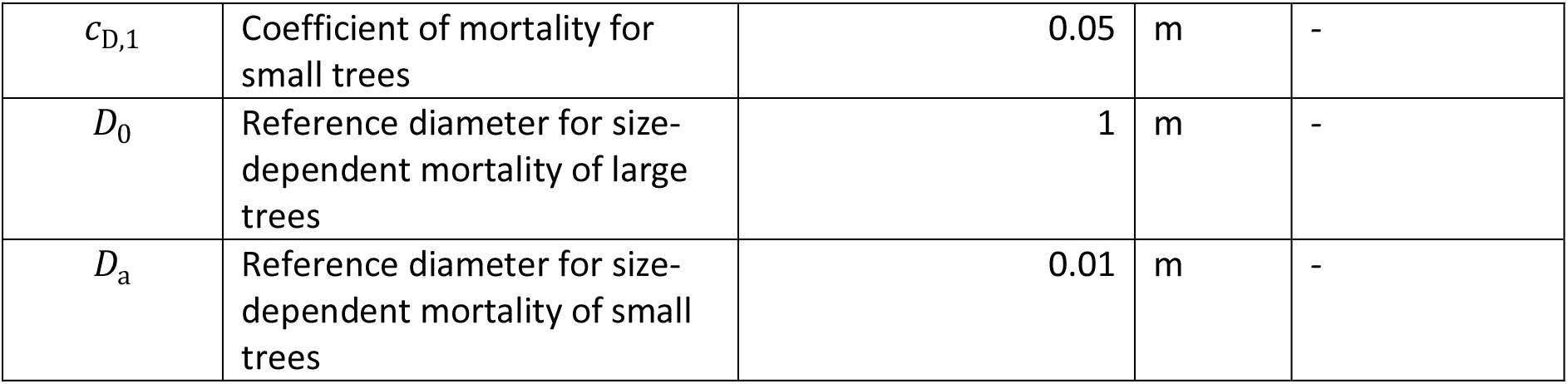
Model traits and parameters. References: Tallo database^79^; GrooT database^80^; Li et al. (2014)^50^; Wenk et al. (2015)^81^; Farquhar et al. (1980)^82^; Doughty et al. (2017)^83^; Camac et al. (2018)^84^.

### Community trait evolution

In the long term, the overall trait distribution of the community evolves by competitive exclusion and directional selection. The strength of competitive exclusion depends on the magnitude of trait-mediated differences in species demographic rates, whereas that of directional selection depends on intraspecific variation. Although we consider light as the only limiting resource, competition for light also implicitly leads to more complex forms of competitive exclusion (e.g., competition for space) due to trait differences in species and emerging population-environment feedbacks. We capture the aggregate effect of trait evolution in terms of changes in community-weighted mean (CWM) traits, defined here as the average of species trait values weighted by the corresponding basal areas of tree stems. As species relative abundances (and basal areas) change due to competition and species traits change by directional selection, the traits of the fittest species are reflected in the CWM trait values. Plant-FATE accounts for competitive exclusion via demographic dynamics and directional selection via evolutionary dynamics theory. In this work, however, we focus on competitive exclusion and suppress directional selection by setting intraspecific trait variance to zero.

## 3 Results

First, we show that the Plant-FATE model reproduces observed forest structure and function in the Amazon Forest. We then use the model to predict ecosystem responses to elevated CO_2_ concentrations.

### 3.1 Model predictions match observations

Forced with current climatic conditions for 3000 years (white regions in Fig. 1), our model’s predictions of emergent ecosystem properties (black lines in Fig. 1; blue curves in Fig. 2) are consistent with observations (grey bands in Fig. 1; grey points and curves in Fig. 2; see Table 2 for sources) across multiple variables characterizing forest structure and function. We quantify forest function in terms of gross and net primary productivities (GPP and NPP, Fig. 1A), canopy stomatal conductance (*g*_c_, Fig. 1B), and average photosynthetic capacity (*V*_cmax_, Fig. 1C).We quantify forest structure in terms of leaf area index (LAI, Fig. 1D), basal area (BA, Fig. 1E), the vertical distribution of leaves in the canopy (Fig. 1F, Fig. S1A), the size distribution of trees (Fig. 2A; with size measured in terms of diameter at breast height, DBH), and the distributions of functional traits (Fig. 2B-D).

**Fig. 1.**
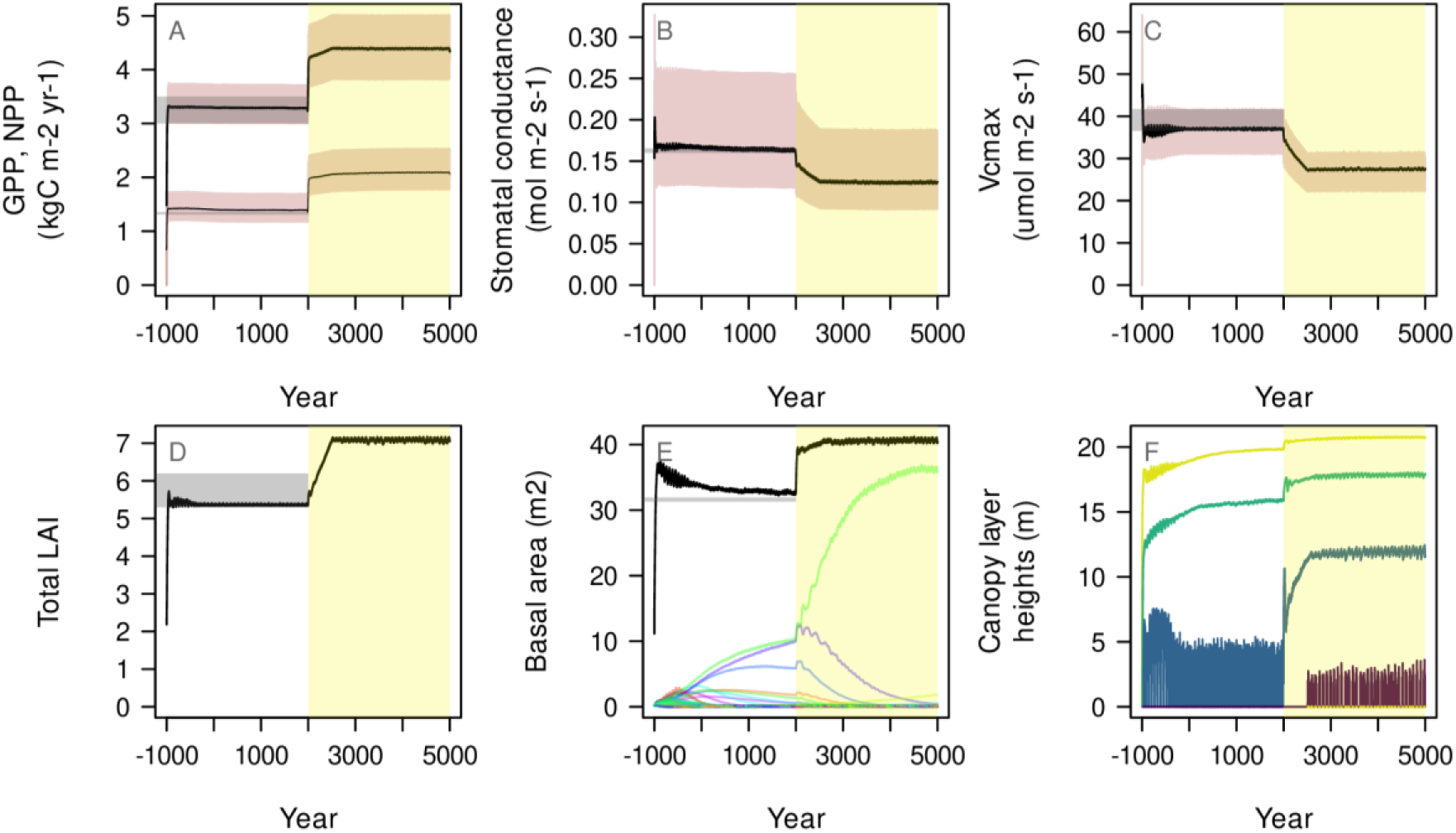
Plant-FATE correctly predicts emergent ecosystem properties under ambient and elevated CO_2_ concentrations. This figure shows model predictions for the Amazon forest site over 6000 years. The first 3000 years (1000 BCE – 2000 CE) constitute a spin-up run under a fixed CO_2_ concentration of 414 ppm (white region in panels A-F), and are followed by 3000 years (2001-5000 CE) with an elevated CO_2_ concentration of 614 ppm (yellow region in panels A-F). Under ambient CO_2_ conditions, predicted (black lines) gross primary productivity (GPP, thick line in panel A), net primary productivity (NPP, thin line in panel A), stomatal conductance per unit ground area (B), photosynthetic capacity per unit crown area (C), leaf area index (D), and total basal area (E; coloured lines show species basal areas) lie within their respective observed ranges (grey horizontal bands; see Table 2 for sources). Canopy closure heights of different canopy layers are shown in (F). Under elevated CO_2_, GPP, NPP, LAI, and basal area increase, whereas stomatal conductance and photosynthetic capacity decrease. In panels A-C, black lines show loess-filtered time series for ease of viewing, whereas brown bands represent the actual time series.

**Fig. 2.**
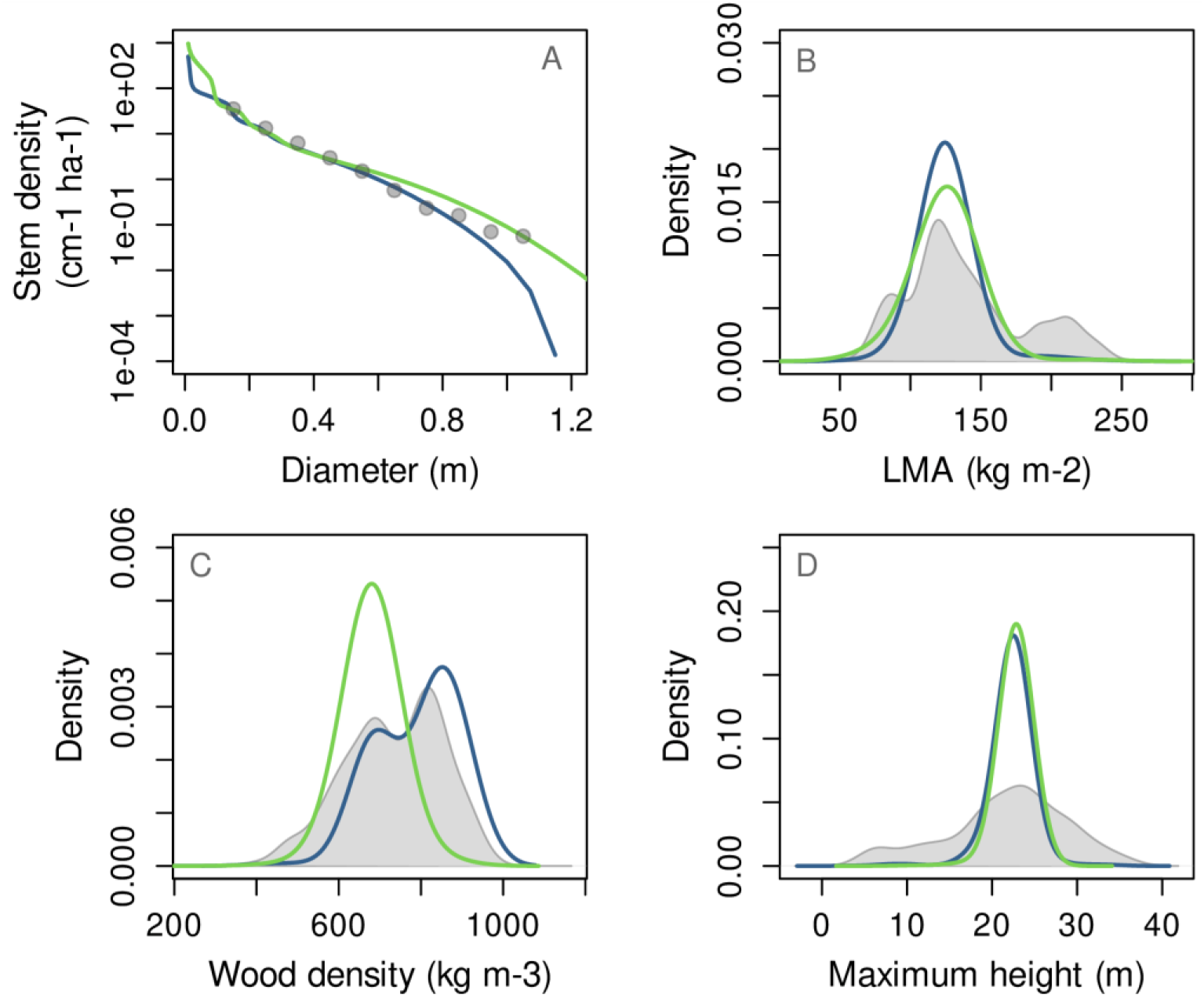
Elevated CO_2_ concentrations drive a community shift towards species with lower wood density. Under ambient CO_2_ concentrations (blue curves), model-predicted equilibrium size distribution (A) and trait distributions (B-D) match those observed at our study site (grey points in A and filled grey curves in B-D). Under elevated CO_2_ concentrations (green curves), our model predicts a shift towards a denser stand with a higher proportion of large trees (A), and a shift towards species with lower wood densities (C).

**Table 2.** Sources for observed values of emergent ecosystem properties used for model calibration and validation. References: Fleischer et al. 2019^11^, Menezes et al. (2022)^85^, Cordeiro et al. (2019)^86^, Vieira et al. (2004)^87^

| Symbol | Ecosystem property | Source |
| --- | --- | --- |
| GPP | Gross primary productivity | Mean annual GPP from K34 eddy-covariance tower, measured between 1999-2013. PI's: C. von Randow et al. (Fleischer et al. 2019) |
| NPP | Net primary productivity | Based on total production in leaves, wood, fine roots and reproduction from two AmazonFACE plots. PI's: I. Pereira, F. Hofhansl, R. Norby, A. Longhi Cordeiro, E. Oblitas et al. (Fleischer et al. 2019) |
| LAI | Leaf area index | Calculated using hemispherical photographs from 2 AmazonFACE plots taken within 3-month intervals during 2015-2017. PI's: F. Hofhansl, S. Garcia, L. Fuchslueger, O. Valverde, K. Schaap, A. Grandis et al. (Fleischer et al. 2019) |
| $V_{\text{cmax}}$ | Average crown-specific photosynthetic capacity | Leaf-specific $V_{\text{cmax}}$ is calculated from measurements at leaf age equalling half the mean leaf lifespan from upper-canopy leaves, as reported in Menezes et al. (2022). This is converted to crown-specific $V_{\text{cmax}}$ by multiplying by crown LAI (= 1.8) and upscaled to a vertical average value by multiplying by the ratio of average light intensity to top-canopy light intensity ($\approx 0.5$). |
| $g_s$ | Stomatal conductance | Measurements from Amazon FACE plots (Menezes et al. 2022) |
| BA | Total stem basal area | Measurements from Amazon-FACE plots |
| AGB | Aboveground biomass | Computed from DBH and height measurements in 8 AmazonFACE plots during 2017 (Fleischer et al. 2019) |
| CFR | Mass of carbon in fine roots | Measurements from 0-90 cm depth range in the AmazonFACE plots (Cordeiro et al. 2019) |
| $u(s)$ | Size distribution | Measurements from 3 plots from Manaus (Vieira et al. 2004) |

Representing an advance over most DGVMs, accounting for trait-size structure allows us to predict the dynamics of size and trait distributions of the community. Equilibrium size distribution predicted by our model under ambient CO_2_ conditions (blue curve in Fig. 2A) matches the corresponding observed distribution (grey points). We obtain predicted and observed community-weighted trait distributions by fitting kernel-density functions to species-specific trait values weighted by species-specific basal areas. For constructing observed trait distributions, we use trait values and basal areas of all species recorded at the study site for which data were available, whereas for constructing predicted trait distributions, we use traits and basal areas of surviving species in model runs. We find that predicted equilibrium trait distributions (blue curves in Fig. 2B-D) match observations (filled grey curves). Since distributions of LMA and maximum height were not used for model calibration, this match provides an independent validation of our model.

Model-predicted aboveground biomass (AGB) is higher than the site-derived value (Fig. S1). However, the model correctly predicts basal area, maximum height, and size distribution, i.e., the three quantities that determine AGB, and for which direct measurements are available. By contrast, the site-derived value of AGB is not a direct measurement, but an estimate based on allometric scaling relationships. Therefore, the mismatch in AGB could also reflect biases or uncertainties in the underlying allometric relationships.

### 3.2 Elevated CO2 concentrations drive a community shift

Starting with the end of the spin-up phase (year 2000 CE), we force the model with elevated CO_2_ concentrations (hereafter called eCO_2_) for another 3000 years (yellow regions in Fig. 1). In this section, we assume that there is no constraint on growth due to nutrient availability (e.g., as a result of external nutrient inputs), and consequently, belowground allocation does not change under elevated CO_2_. Meteorological variables (temperature, VPD, PAR, and soil water potential) are recycled from the 2000-2015 period as in the spin-up run. This arrangement allows us to isolate the effect of eCO_2_ alone, without the confounding effects of changes in plant strategies (such as allocation patterns) or in other meteorological variables (such as temperature).

For a 200 ppm (∼48%) increase in CO_2_ and in the absence of nutrient limitation, our model predicts a 33% increase in gross primary productivity (GPP), despite a reduction in photosynthetic capacity (−25.8%) and stomatal conductance (−23.6%). Autotrophic respiration increases by just 8%, leading to a 49.9% increase in net primary productivity (NPP). The equilibrium size distribution under eCO_2_ (green curve in Fig. 2A) has a higher amplitude and fatter tail compared to that under ambient CO_2_ (blue curve). Thus, eCO_2_ leads to an increase in the total number of trees as well as an increase in the proportion of large trees. Overall, this results in ∼24.5% increase in basal area and ∼15.8% increase in aboveground biomass (AGB). The increase in standing biomass is less than the increase in productivity due to a shift in species composition towards fast-growing (acquisitive) species with lower wood density, as explained in further detail below.

Elevated CO_2_ leads to an increase in community leaf area index, and thus an increase in the number of fully filled canopy layers (from three under ambient CO_2_ to four under eCO_2_; Fig. 1F). The bottom edge of the third layer rises from ∼2m to ∼12m, which decreases light availability in the understorey. This triggers a shift in community composition towards species that can grow quickly and escape the dark understorey, such as those having lower wood densities (green curve in Fig. 2C). This is reflected in a community shift towards species with low wood density compared to that under ambient conditions (blue curve in Fig. 2C). However, species with lower wood density experience increased mortality rates, leading to faster turnover (lower residence time) of trees in the ecosystem. This combination of a decrease in wood density and an increase in biomass turnover dampens the increase in standing biomass relative to a corresponding increase in GPP.

### 3.3 Population-environment feedbacks determine the direction of the community shift

Next, to confirm our conclusion that darkening of the understorey is the cause of the emergent community-level response, we focus on the performance of individual trees growing in the light environment set by the community. The optimal strategy for individual trees is the one that maximizes their fitness, defined in terms of the lifetime reproductive output. We calculate the fitness *F* of trees with different wood densities as follows,

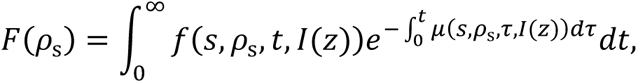

with

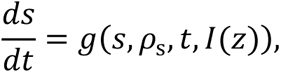

where *g*(⋅), *μ*(⋅), and *f*(⋅) are the growth, mortality, and fecundity rates as a function of the plant size *s*, wood density *ρ*_s_, time *t*, and the vertical light profile *I*(*z*). The integrand in the above equation is the expected seed output at time *t*, which is a product of the fecundity of the plant at time *t* and the probability that the plant survives until time *t*.

We calculate fitness landscapes *F*(*ρ*_s_) under a fixed vertical light profile *I*(*z*), i.e., the focal plant does not affect the light environment, but it may experience different light levels as it grows. We first set the vertical light profile to that predicted by the full model under ambient CO_2_ concentrations (corresponding to year 2000 CE in Fig. 1F). In this light environment, the fitness landscape is hump-shaped (Fig. 3A), because trees with low wood densities have high mortality, whereas trees with high wood density grow too slowly. Under ambient CO_2_ concentrations, the fitness peaks at a wood density of ∼800 kg m^-3^ on average, whereas under eCO_2_, it peaks at a higher wood density of over 1200 kg m^-3^. This implies that elevated CO_2_ should favour trees with higher wood density, as long as the light environment remains unchanged. However, when we additionally set the light profile to that predicted from the full model under elevated CO_2_ concentrations (corresponding to year 5000 CE in Fig. 1F), the fitness peak shifts to lower wood densities (∼650 kg m^-3^).

**Fig. 3.**
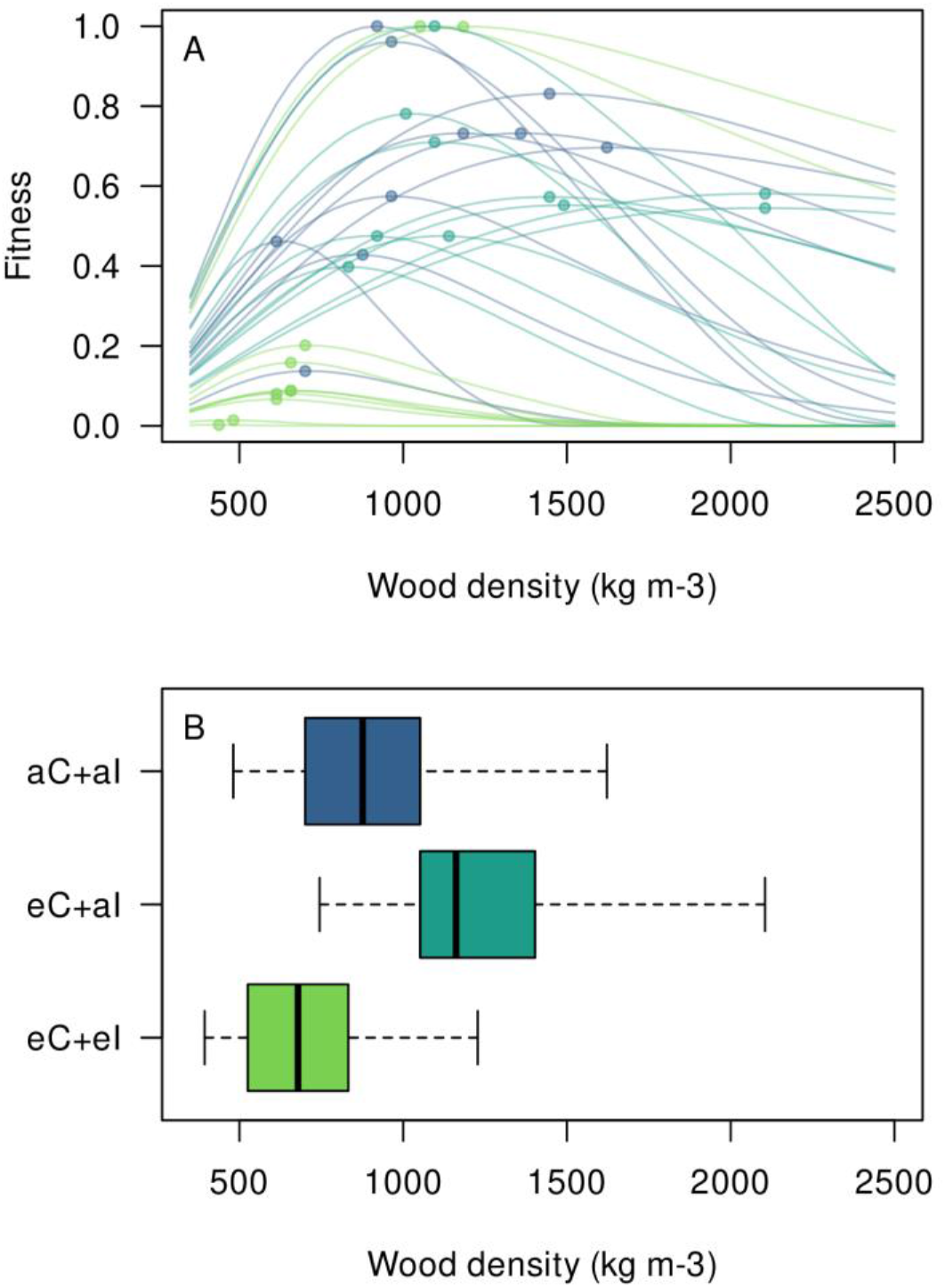
In a community of trees, wood density evolves in a direction opposite to that in isolated trees. (A) Fitness of individual trees growing in a specified light environment, as a function of wood density. Each curve corresponds to a family of species with different wood densities, but with identical values of LMA, maximum height, and *ψ*_50x_. Thus, each curve shows the prospective fitness of a species *if* it had a given wood density, all other traits being equal. Blue curves are obtained with ambient CO_2_ and ambient light environment (aC+aI), dark green curves are obtained under elevated CO_2_ but ambient light environment (eC+aI), whereas light green curves are obtained under elevated CO_2_ and an altered light environment as observed under elevated CO_2_ (eC+eI). For all species and scenarios, fitness peaks at intermediate values of wood density (marked by points), reflecting the trade-off between growth and survival. (B) Optimal wood density (corresponding to the fitness maxima in panel A) is higher in the eC+aI scenario compared to the aC+aI (baseline) scenario, implying that under elevated CO_2_ but in the absence of environmental feedbacks, trees with higher wood density are fitter. However, when environmental feedbacks are accounted for (eC+eI scenario), optimal wood densities are lower than the baseline.

Thus, the effects of eCO_2_ on a tree community may be opposite to those on isolated plants. This is in line with evidence from controlled experiments on individual plants versus those on in-situ forest stands^52^. For isolated trees, eCO_2_ allows for faster growth, and therefore trees benefit from higher wood densities that confer lower mortality rates. In a community of competing trees, however, an increase in leaf area under eCO_2_ intensifies competition for light. In order to survive, juvenile trees need to grow faster to escape the dark understorey and emerge into successively higher and brighter layers. This enhances the survival of fast-growing trees with lower wood density, triggering a community shift towards such a strategy.

### 3.4 Increased belowground allocation dampens aboveground carbon-sink strength

Next, we analyse the effect of nutrient availability on the community responses to elevated CO_2_. A majority of Free Air Carbon Enrichment (FACE) experiments have shown that, under eCO_2_, trees allocate relatively more carbon below ground into nutrient-acquiring mechanisms, including fine roots, root exudates, and mycorrhizae^12, 53–55^. Such investment alleviates nutrient limitation in infertile soils, allowing trees to make better use of increased CO_2_ availability. Currently, Plant-FATE does not have an explicit representation of nutrient cycling, and thus cannot predict the magnitude of change in allocation patterns. However, it is possible to explore how different levels of allocation shifts affect overall community-level forest responses. In Plant-FATE, the allocation strategy is reflected in the ratio of fine-root mass to leaf area, *ζ*, but since we do not resolve belowground C pools, *ζ* may also be interpreted as the total investment in fine roots, root exudates, and mycorrhizae. Under ambient CO_2_ conditions, we calibrated the value of *ζ* = 0.2 kg m^-2^ to obtain the correct value of fine root mass (Fig. S1). In this section, we simulate three eCO_2_ scenarios in which belowground C allocation increases by 25%, 50%, or 100% under eCO_2_.

First, we predict that allocation shifts strongly dampen the eCO_2_-driven increase in leaf area. Even a 25% increase in ζ is sufficient to negate the potential increase in LAI (Fig. 4C). Second, since the fraction of absorbed radiation saturates with increasing LAI, changes in the LAI of a closed-canopy forest have only a small effect on GPP, leading to a relatively insensitive response of GPP to allocation changes (Fig. 4B). Third, however, higher belowground investment implies higher fine-root respiration, so that NPP responds strongly to allocation shifts. A 100% increase in fine-root allocation completely reverses any increases in NPP by CO_2_ fertilization (Fig. 4D). Fourth, consequently, increases in AGB are dampened by allocation shifts to relatively higher belowground biomass, and completely reversed by a 100% increase in belowground allocation. Finally, the magnitude of the increase in LAI also affects the magnitude of the community shift under eCO_2_ – while a 25% increase in fine-root allocation still leads to a (less intense) community shift, a 50% increase prevents it entirely. Since a 100% increase in belowground allocation completely negates any increase in aboveground leaf area and biomass, this is a limit beyond which it becomes detrimental for trees to invest more carbon belowground.

**Fig. 4.**
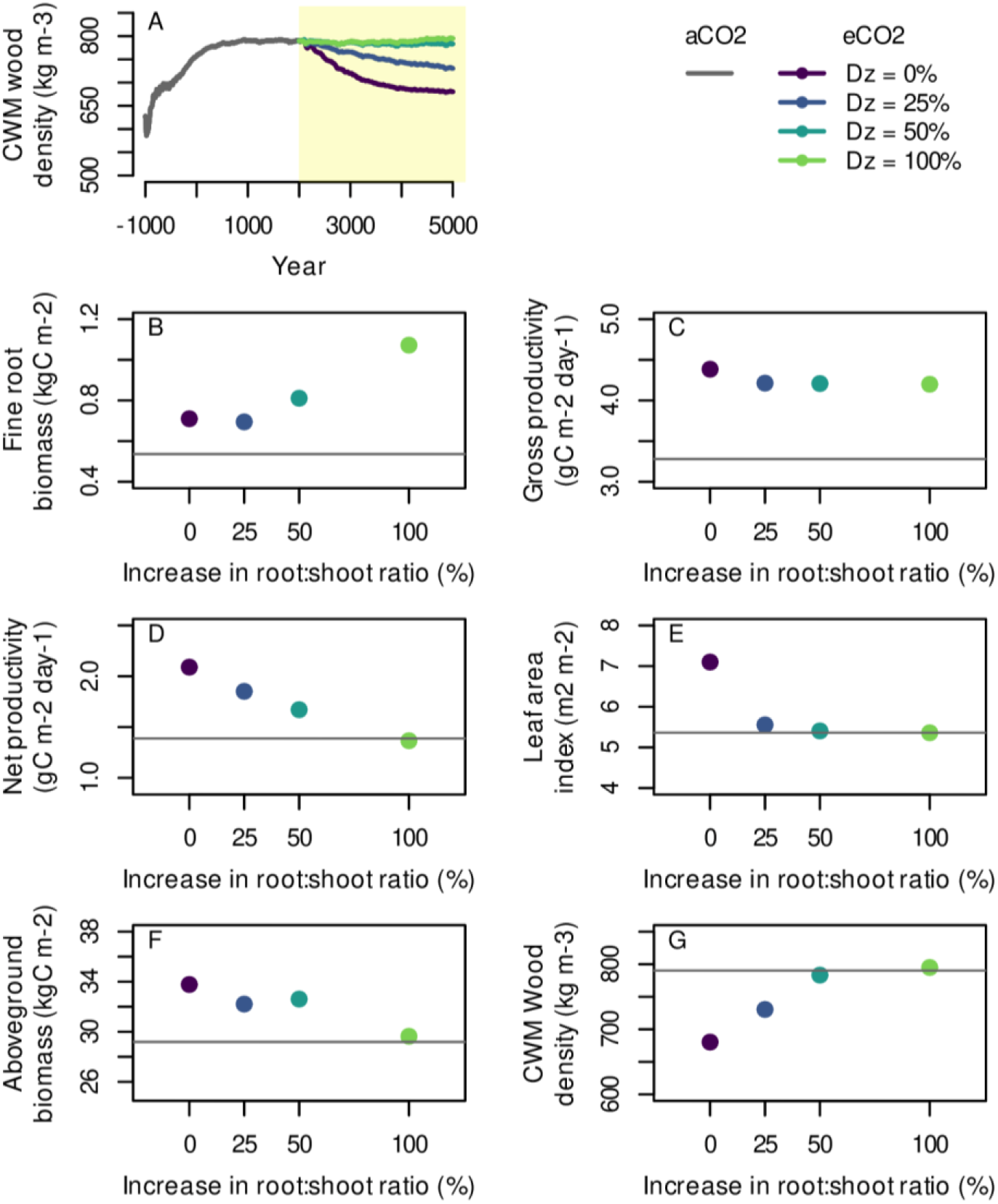
Shifts in allocation towards greater belowground investments limit the increase in aboveground biomass but also stabilize the community. (A) Time series of community-weighted mean wood density for different levels of allocation shifts. (B-G) Equilibrium emergent ecosystem properties under elevated CO_2_ concentrations for different levels of allocation shifts (coloured points) compared to the corresponding value under ambient CO_2_ (horizontal line).

**Fig. 5.**
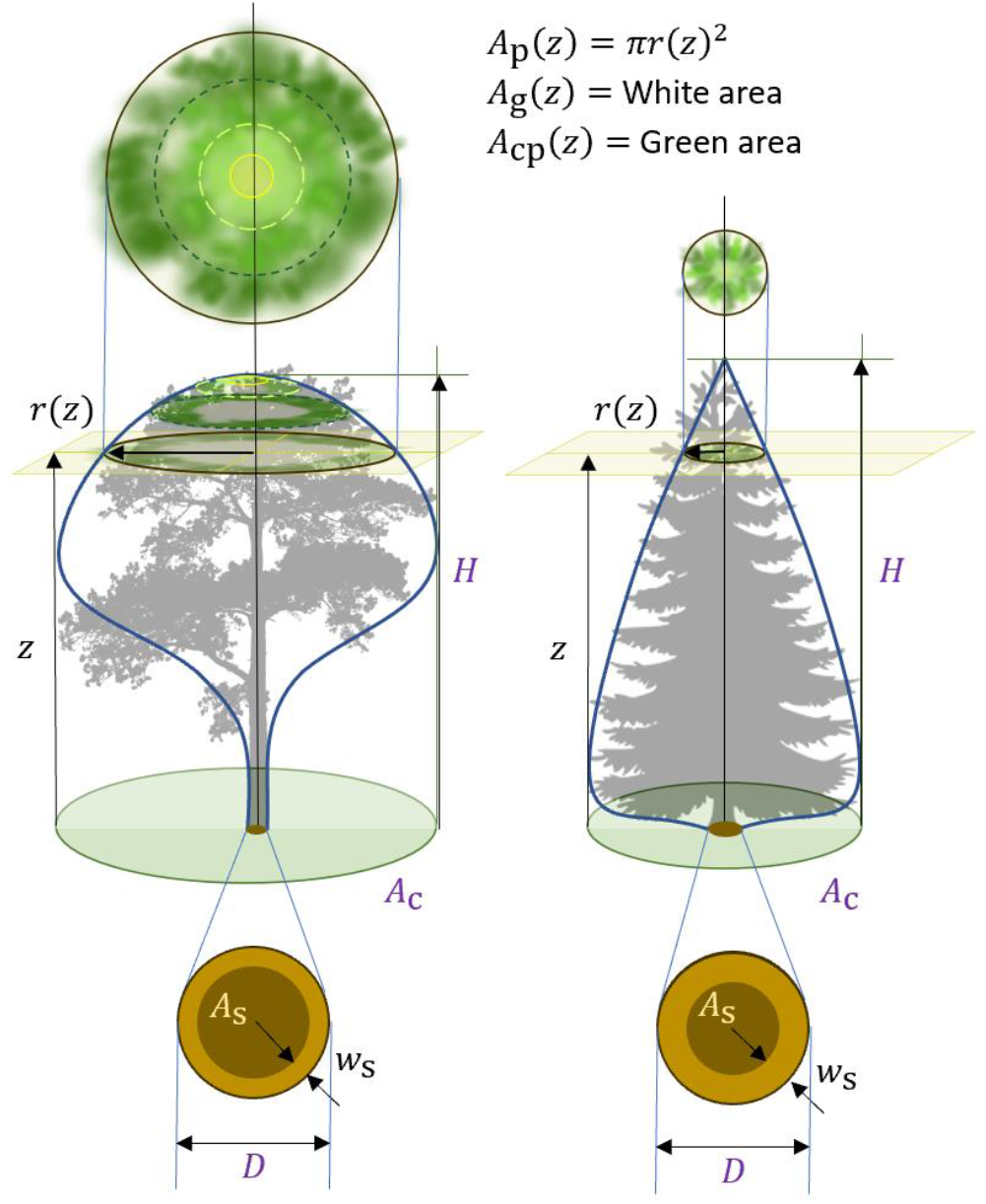
Tree architecture in Plant-FATE. Tree size is defined by the basal diameter *D*, from which projected crown area *A*_c_, sapwood area *A*_s_, sapwood width *w*_s_, and height *H* are calculated using dimensional scaling relationships. Trees have a vertically explicit crown, whose shape is given by a function *r*(*z*).

## 4 Discussion

We set out to understand the multiscale adaptive responses of biodiverse forests to elevated CO_2_. To that end, we have developed a new trait-size structured eco-evolutionary vegetation model, Plant-FATE, that accounts for the physiological acclimation of plastic traits, demographic dynamics of species abundances, and the evolutionary dynamics of genetic traits. By drawing upon key emerging concepts and theories in vegetation modelling, such as eco-evolutionary optimality principles, trait-size-structured vegetation demographics, and evolutionary dynamics theory, our model correctly predicts the structure and function of a hyper-diverse central-eastern Amazonian terra-firme forest across multiple emergent ecosystem properties. We have explored three key aspects of ecosystem responses to elevated CO_2_. First, we have identified three mechanisms operating at three distinct temporal and organizational scales: short-term plant-level responses, e.g., in terms of photosynthetic acclimation, mid-term demographic responses in terms of changes in species abundances, and long-term structuring of the community by competitive exclusion. Second, we have shown that under elevated CO_2_ and nutrient-fertilized soils, productivity, leaf area, and aboveground biomass increase. Such fertilization concomitantly intensifies competition for light and triggers a shift in community composition towards fast-growing but fragile species characterized by lower wood densities and higher mortality rates. Third, in infertile soils, eCO_2_-driven increases in belowground carbon allocation may dampen the CO_2_ fertilization effect and prevent a competition-induced community shift, allowing the ecosystem to remain in a steady-state (Fig. 4A).

### 4.1 Comparison with observations from FACE experiments

Free Air Carbon Enrichment (FACE) experiments provide valuable information on the multidimensional response of forests to elevated CO_2_. Since data from the Amazon FACE experiment is not yet available, we compare our model predictions with observations from a similar experiment conducted in Australia (EucFACE). Although EucFACE is located in a different forest type (savanna) compared to the Amazon (wet evergreen forest), it shares some similarities with Amazon FACE, i.e., it is conducted in a mature forest with phosphorus-limited soils. Therefore, EucFACE provides a suitable reference point for comparing our model results, which we offer as prior hypotheses for the Amazon FACE experiment.

At the EucFACE site, eCO_2_ increases GPP, but only a small fraction of the carbon surplus ends up in ecosystem carbon pools. Instead, most of the additional carbon is respired back into the atmosphere via increased soil respiration^56^. Furthermore, there is no change in the biomass of fine roots and mycorrhizae, suggesting that the extra carbon probably enters the soil via root exudates or increased fine-root turnover, before being respired back into the atmosphere. The role of nutrient limitation in these experiments is also confirmed by nutrient fertilization experiments which show an increase in net primary productivity upon phosphorus addition^57, 58^.

Our model results are consistent with these observations, particularly when an allocation shift is considered. Specifically, with a doubling of belowground investment, we predict that a 48% increase in CO_2_ concentration elicits a 28.5% increase in GPP, which is entirely lost via respiration, leading to no increase in NPP, leaf area, or aboveground biomass. Although the increased respiration comes from an increase in fine root biomass in our model setup, an increase in the production of root exudates – rather than fine root biomass – would lead to a similar outcome. Explicit modelling of root exudates would allow us to resolve such belowground dynamics and would thus be a promising model extension.

The EucFACE experiment falsified and rejected three common hypotheses that vegetation models often use to predict eCO_2_ effects on forests. These are: (1) eCO2 increases GPP, which in turn proportionally increases biomass; (2) nutrient availability downregulates the eCO_2_-driven increase in GPP; or (3) enhanced productivity under eCO2 is simply lost by aboveground respiration. Our model also does not support these hypotheses. Rather, our model supports the EucFACE findings that eCO_2_ leads to increased GPP, but the surplus carbon goes into the soil rather than into aboveground biomass, from where it is respired back into the atmosphere.

We also compare our model results with a meta-analysis of FACE experiments. First, our model predictions of an increase in GPP accompanied by a decrease in photosynthetic capacity and stomatal conductance are consistent with a consensus of FACE findings^59^. Another key observation from FACE experiments is that eCO_2_ enhances canopy leaf area in forests with low LAI, but not as much in forests with high (>5) LAI. At the same time, nutrient fertilization experiments conducted on saplings in the Amazon^60^ have shown an increase in leaf area (although it remains unclear whether mature trees would show a similar trend). Our model predictions are consistent with these observations: we predict an increase in LAI when there is no increase in belowground allocation under eCO_2_ (this is reminiscent of a nutrient fertilization scenario), especially due to an increase in tree density in the understorey. However, even a slight (25%) allocation shift to nutrient acquisition prevents an increase in LAI. This is because the fraction of absorbed light saturates with increasing LAI, and an increase in LAI beyond 5 brings diminishing returns in terms of carbon uptake. One key difference between observations and our model predictions is that experiments show an increase in leaf area within individual trees, whereas our model-predicted increase in LAI results from an increase in total crown cover. This is because we do not treat crown LAI as a plastic trait. Extending our model by accounting for individual-level acclimation of crown LAI would be a promising way forward.

### 4.2 Comparison with other trait-based and demographic models

In recent years, a new generation of trait-based vegetation models has emerged (see ref.^20^ for a comprehensive review). The key strengths of Plant-FATE compared to many other models are (i) a multidimensional representation of functional diversity, accounting for inter- and intra-specific trait variation combined with phenotypic plasticity, (ii) the use of EEO principles for deriving parameter-sparse yet more robust models of plastic trait acclimation (especially for modelling the acclimation of photosynthesis and leaf lifespans in the present study), and (iii) an ecologically realistic yet theoretically and computationally efficient approach to unify physiological, demographic, and evolutionary processes that determine ecosystem function. As a result, Plant-FATE can predict emergent ecosystem properties across multiple structural and functional target variables with fewer parameters and processes compared to many other trait-based demographic models.

A major strength of trait-based models is that they can predict community assembly under given environmental conditions, or changes in community composition in response to environmental changes. We compare our model predictions with those of other similar models. E.g., a study with the aDGVM model^47^ showed how fire suppression causes a community shift towards species with greater fine-root allocation and higher wood density. Similarly, a recent study with the CAETE model^61^ predicted that drought induces a reorganization of the community towards species that allocate greater carbon to fine roots to improve access to water, but at the expense of a decrease in aboveground productivity. Our model also predicts a community shift in response to environmental change (eCO_2_) that is subject to a trade-off in aboveground vs belowground investment. However, we show explicitly how height-structured competition for light critically affects the selection pressures on species, leading to counterintuitive changes in the direction of the community shift.

### 4.3 Model limitations and possible extensions

An important limitation of our model is that it currently does not simulate closed water and nutrient cycles. Thus, although we could explore the consequences of different levels of allocation shifts, we could not predict exactly how much of a shift we should expect under elevated CO_2_. Closing the mass balance for water and nutrients and modelling nutrient acquisition dynamics will allow us to consider the costs and benefits of increased belowground investment, enabling prediction of the best allocation strategy under ambient and elevated CO_2_ conditions.

Currently, we have incorporated only the most essential eco-physiological processes into our model with a bare minimum number of parameters. While these are sufficient to represent a wet evergreen forest like the Amazon, the results presented here are of an exploratory nature, analysing simple what-if scenarios. Indeed, despite its simplicity, our model seems to correctly capture emergent structural and functional properties of a hyperdiverse tropical forest. Nonetheless, making the model applicable to a wider range of biomes will require incorporating additional features, such as seasonal phenology, belowground hydraulics, reproductive trade-offs, and flexible allometric scaling patterns. Each of these model extensions is a promising direction for future work.

### 4.4 Adaptive processes and their timescales

Our work reveals three broad timescales over which forests respond to climate change. First, the time series of GPP, stomatal conductance, and *V*_cmax_(Fig. 1) show that a step increase in CO_2_ in 2000 CE elicits an almost instantaneous response in these variables, which occurs on the weekly timescale corresponding to photosynthetic acclimation. The same time series show a further change in these variables over a timescale of about 500 years. This is due to a gradual change in the demographic distribution of the community and the vertical light profile. This timescale corresponds to that on which the community structure reaches a new demographic equilibrium (Fig. 1F). Finally, Fig. 3 shows that the genetic trait distribution (in our model runs, the wood density distribution) equilibrates on a timescale of about 2000 years. This is the timescale over which community shifts resulting from competitive exclusion could occur. While these three timescales are broadly illustrative of the three processes (physiological acclimation, demographic equilibration, and competitive exclusion) in a community of long-lived trees, changes in specific traits could occur much faster due to phenotypic plasticity. For example, in our model, crown LAI and wood density of individual trees are held constant throughout their lifetime and thus change only via changes in species composition. In reality, however, these traits have sufficient plasticity to change over annual timescales through new leaves and wood. Therefore, in real forests, LAI and wood-density changes could occur faster (on the respective acclimation timescales) than reported here.

### 4.5 Adaptive responses beyond the tropics

The adaptive responses identified in our modelling study readily extend to similar observations beyond the tropics. For example, studies conducted in temperate ecosystems comprising tree species such as spruce, pine, and oak have reported a decline in tree wood density over the past century. Wood density is also reported to decrease with nitrogen fertilization^17^. Furthermore, stand thinning may also alter wood density of conifers, but these changes are confounded by a decrease in stem slenderness^62^. The mechanisms behind these changes remain unclear^18, 19^. It has been hypothesized that the observed trends over the past century could result from increasing anthropogenic nitrogen deposition, prolonged growing season lengths, and higher temperatures^19^. Based on our model, we propose a new mechanism for the observed trends – competition for light. Our model predicts that any factor that increases leaf area (e.g., CO_2_ or nutrient fertilization or an increase in stand density) can trigger environmental feedbacks that drive a decrease in wood density (and likely an increase in stem slenderness, but we did not test this in the present study). One difference between the observed trends and our model-based predictions is that the observed trends are due to gradual phenotypic changes in the same individuals over time, whereas the model-predicted trends are due to a community shift. Thus, although the model correctly captures the trend in wood density, it predicts a much longer timescale for this change. This mismatch can be readily resolved by accounting for the annual-scale acclimation of wood density in our model, and doing so would be a promising avenue for future model development.

### 4.6 Ecosystem-level responses to climate change

Our work highlights the importance of competition for resources in modulating community-level dynamics of ecosystems. We have shown that selection pressures on trees competing for light depend on the relative abundance of individuals (i.e., they are frequency-dependent), and can be different from (and indeed, opposite to) those on isolated trees. Therefore, feedbacks between the population and the environment resulting from species interactions should also be considered while designing experiments aimed at assessing forest responses to eCO_2_. Experiments that expose individual plants to eCO_2_ may yield responses that do not hold up on the level of communities. We therefore also recommend caution when using meta-analyses of single-species experiments to predict global responses to climate change. Instead, experiments on the community level, such as the FACE experiments, are better suited to assess ecosystem-level responses.

One of the extreme consequences of frequency-dependent selection is evolutionary suicide, where evolution drives a viable population to extinction^63^. We find a glimpse of this effect in our study. Elevated CO_2_ and nutrient concentrations select for fast-growing trees that attempt to maximize their self-interest by growing taller than their neighbours, but this happens at the expense of reduced wood density. A competition-induced reduction in wood density may render global forests more vulnerable to climatic extreme events^64, 65^ that are expected to occur at higher frequencies in the future.

## 5 Acknowledgements

We thank the following researchers for assisting with the compilation of data and making it available for model calibration and validation: Celso von Randow, Alessandro Araujo, and Katrin Fleischer (meteorological data from K34 eddy covariance flux tower, collected by the LBA Program: The Large Scale Biosphere-Atmosphere Research Program in the Amazon); Lucia Fuchslueger and Karst Schaap (volumetric soil water content); Juliane Menezes (stomatal conductance); and Cleiton Eller and Gabriela Sophia (xylem P50). We also thank Jeremy Lichstein for feedback on the implementation of the PPA within our model. JJ and UD acknowledge funding from the European Union’s Horizon 2020 research and innovation programme under the Marie Skłodowska-Curie Actions Individual Fellowship (grant agreement No. 841283 – Plant-FATE). JJ and FH acknowledge funding from the Strategic Initiatives Program of the International Institute for Applied Systems Analysis (IIASA; Project name – RESIST). UD acknowledges funding from the Global Bioconvergence Center of Innovation at the Okinawa Institute of Science and Technology Graduate University, OIST, supported by a grant from the Japan Science and Technology Agency, JST, Program for an Open Innovation Platform for Academia-Industry Co-Creation, COI-NEXT. JJ, FH, OF, ÅB, and UD acknowledge funding from IIASA and the National Member Organizations that support the institute.

## 6 Author contributions

JJ developed the Plant-FATE model with inputs from FH, ICP, and UD. JJ, BDS, and FH designed the study. OF, ÅB, and UD contributed to theoretical development and/or provided complementary analysis tools. JJ and SS calibrated the model and performed the model runs. CCB, IFA, and DL provided data from the AmazonFACE site. JJ developed the software package and wrote the first draft of the manuscript. All authors contributed to revised versions of the manuscript.

## 7 Methods

In this section, we briefly describe the formulations of key processes in the Plant-FATE model. A full derivation of all model components can be found in the Supplementary Information.

### 7.1 Physiology

**Photosynthesis, respiration, and turnover.** Gross photosynthesis rate per unit crown area (*P*_c_) is computed using the P-hydro model^28^, which predicts the simultaneous acclimation of three plastic traits: two photosynthetic capacities (*V*_cmax_ and *J*_max_) and stomatal conductance (*g*_c_). To do so, leaves maximize profit (Φ) on a weekly timescale, defined as the benefit from assimilation minus the costs arising from maintaining the photosynthetic machinery and the risks of hydraulic failure,

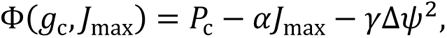

where *α* and *γ* are unit costs, and Δ*ψ* is the soil-to-leaf water-potential difference that depends on whole-plant hydraulic traits such as conductivity *K* and vulnerability *ψ*_50_.

Net whole-plant photosynthesis is calculated as

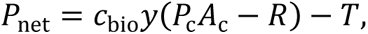

where *c*_bio_ is the biomass produced per mol CO_2_, *y* is the yield factor accounting for carbon lost in growth respiration, *R* is the respiration rate of leaves, fine roots, and sapwood, and *T* is the turnover rate of leaves and fine roots.

Sapwood respiration rate is assumed to be proportional to sapwood mass, whereas fine-root respiration rate is assumed to be proportional to both fine-root mass and gross productivity. Optimal leaf turnover rate (*τ*^∗^) is calculated using the theory of leaf economics^31^, which yields

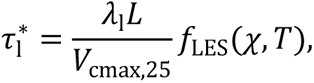

where *λ*_l_ is leaf mass per unit area, *L* is the leaf area index within the tree crown (of an individual tree), and *f*_LES_ is a scalar that depends on the leaf-internal-to-external CO_2_ ratio *χ* and temperature *T*, and incorporates the cost of leaf construction. Fine-root turnover rate is calculated using a similar approach.

### Crown architecture and dimensional scaling

Trees in Plant-FATE have a vertically explicit crown structure (Fig. 5), with crown radius *r*(*z*) at each height *z* given by

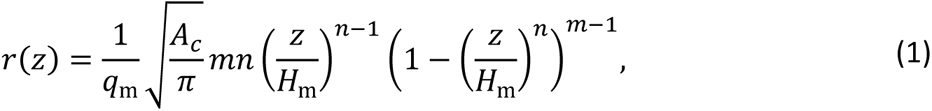

where *H*_m_ is the maximum height for the species, *m* and *n* are traits determining crown shape, *A*_c_ is the projected crown area, and *q*_m_ is calculated so that 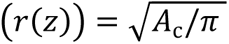. The traits *H*_m_, *m*, *n*, and *A*_c_ are treated as species-specific genetic traits. Sapwood mass is calculated using the generalized pipe model, assuming a constant sapwood-area-to-crown-area ratio throughout the crown.

Tree size is defined in terms of the basal diameter *D*. All other size-related variables (such as height *H*, crown area *A*_c_, fine root mass *m*_r_) are connected to *D* via dimensional scaling relationships. These are derived from an extended version of the T-model^50^, as follows,

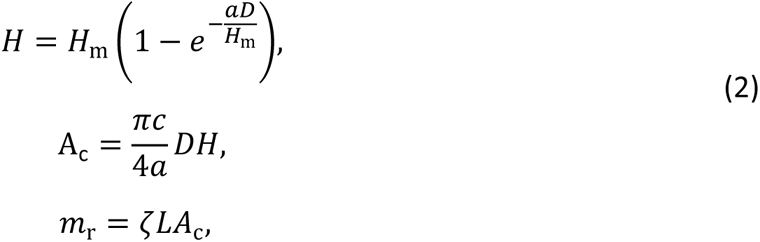

where *a* is the stem slenderness ratio, *c* is the ratio of crown area to sapwood area, and *ζ* is the root-to-shoot ratio. All three traits (*a*, *c*, *ζ*) are treated as genetic traits, but *ζ* is altered to various degrees under eCO_2_.

### 7.2 Demographic dynamics

The dynamics of the size distribution of individuals in the population is given by the McKendrick-von Foerster equation,

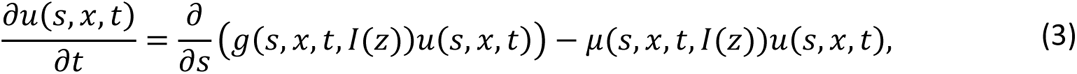

where *u*(*s*, *x*, *t*) is the density distribution of individuals with size *s* and traits *x* at time *t* (expressed as the number of individuals per unit size per unit ground area), *g*, *μ*, and *f* are the growth, mortality, and fecundity rates, respectively, calculated from the biomass allocated to these processes, and *I*(*z*) is the vertical light profile (i.e., the feedback environment) collectively created by all individuals in the population.

The fraction of biomass *f*_r_(*D*) allocated to reproduction is a function of diameter,

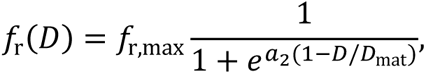

where *f*_r,max_ is the maximum fraction allocated to reproduction, *D*_mat_ is the diameter when the tree reaches 80% of its maximum height, and *a*_2_is a parameter describing the rate at which the plant increases reproductive investment. This leads to a fecundity rate of

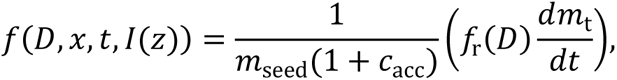

where *m*_seed_ is the seed mass, *c*_acc_ is the cost of seed accessories, and *dm*_t_/*dt* is the net productivity available for growth and reproduction.

The remaining fraction (1 − *f*_r_(*D*)) of production is allocated to growth, yielding a growth rate of

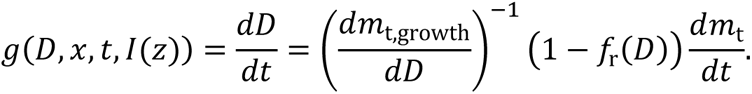

Mortality rate depends on diameter, growth rate, and wood density,

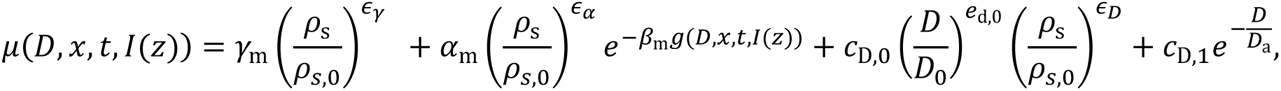

where *ρ*_s,0_, *γ*_m_, *β*_m_, *α*_m_, *c*_D,0_, *c*_D,1_, *D*_0_, *D*_a_, *ϵ*_*γ*_, *ϵ*_*α*_, and *ϵ*_D_ are constants.

The PDE (3) has the following boundary condition which accounts for the recruitment of new individuals,

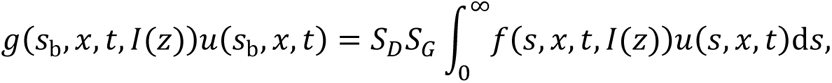

where *s*_b_ is the size at birth, *S*_D_ is the probability of seed survival during dispersal, and *S*_G_ is the probability of sapling survival until recruitment.

### 7.3 Competition for light

We assume that individual trees can optimally place their crowns in canopy gaps to minimize shading from neighbours. Trees can achieve this by slightly bending their stems. This behaviour is called the “perfect plasticity assumption” (PPA)^51, 66^. If all trees place their crowns optimally, the joining heights of crown pairs converge to a single value, *z*^∗^, such that the total crown area above *z*^∗^equals the ground area. Once the canopy closes with one layer of crown, the remaining crowns below this layer do the same, giving a second canopy closure height 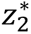, and so on. The heights 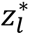 for the *l*^th^ canopy layer can be calculated by solving

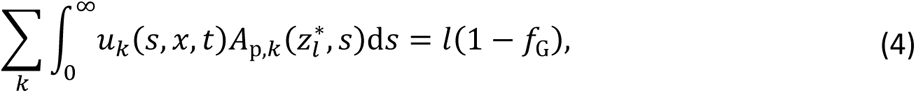

where 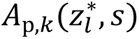 is the projected crown area of a tree of species *k* and size *s*, and *f*_G_ is the fractional gap area between tree crowns (note the difference with *f*_g_, which is the fractional gap area within tree crowns).

The PPA model also allows us to compute light transmission through the canopy layers. The average fraction of light absorbed by layer *l* is given by

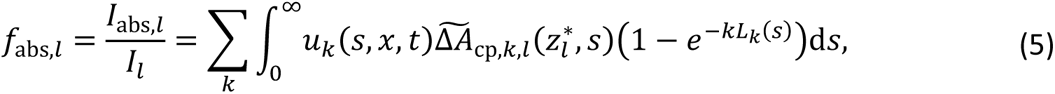

where 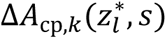 is the filled crown projection area of a tree of species *k* and size *s*, and *L*_*k*_ is the within-crown LAI of species *k*. The average light intensity at the top of layer *l* + 1 is the light intensity transmitted through layer *l*, given by

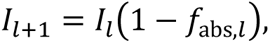

with *I*_1_ being the top-canopy incident light intensity.

### 7.4 Model calibration and validation

We calibrate and test the model at the site scale, using data from the AmazonFACE experimental site located at the Experimental Station of Tropical Forestry ZF2 (2°35’ S; 60°12’ W). Meteorological forcing data (monthly averages of mean air temperature, vapor pressure deficit, daily mean photosynthetic active radiation *PAR*, and daily maximum *PAR*) were obtained from the Amazonas (Manaus) - ZF2 km 34 (K34) eddy covariance flux tower, located approximately 2 km from the AmazonFACE site^67^ (2°36’32.67’’S, 60°12’33.48’’W). Atmospheric CO_2_ concentration values from 2000 to 2020 were calculated by adding +200 to global estimates^68–70^ for these years. Soil water potential was calculated based on site values of volumetric soil water content and soil texture^71, 72^. For calibration, we run the model under a CO_2_ concentration of 414 ppm (corresponding to current conditions in 2020 CE) and forced with meteorological variables from 2000-2015 (repeated periodically).

Data for constructing observed distributions of functional traits were compiled from the following sources: LMA and maximum height: AmazonFACE plots; wood density: Global Wood Density Database^73^; xylem *ψ*_50_: refs.^74–77^. Sources for data on other emergent ecosystem properties used for model calibration and validation are listed in Table 2. Other than functional traits that can be species-specific, Plant-FATE inevitably uses several constant parameters. Wherever possible, we minimize the number of such parameters by utilizing optimality principles. Parameters of EEO-based model components are derived independently (i.e., from sources other than the AmazonFACE site) from their respective global calibrations. Some parameters, e.g., those related to mortality, are derived from the literature. This leaves us with a total of seven unknown parameters which are either difficult to estimate empirically or for which sufficient data is unavailable (Table 1). These parameters are tuned such that model predictions match site observations. Observed distributions of LMA and maximum height are not used for parameter tuning, and thus provide an independent validation of model predictions.

### 7.5 Model runs

We initialize the model with 100 random functional species, i.e., for each species, the four aforementioned traits are drawn from a uniform distribution spanning observed ranges for each trait. All species are initialized with identical size distributions that subsequently change due to demographics and competition. An immigrant species with random traits is introduced into the community every 20 years. Species whose density falls below a certain threshold are removed from the population. The model runs at a fortnightly timestep. Each model run consists of a spin-up phase lasting 3000 years (1000 BCE – 2000 CE). During this phase, values of environmental forcing variables from 2000-2015 are periodically extended and used as model inputs, and CO_2_ is set to a fixed value of 414 ppm (representative of year 2020). With this forcing, the ecosystem reaches a quasi-steady state – quasi because emergent ecosystem properties (such as productivity, leaf area index, basal area) reach a steady state, but relative species abundances may continue to change slowly. After spin-up, the model is forced with an elevated CO_2_ scenario for another 3000 years (2001 CE – 5000 CE), in which CO_2_ concentration increases to 614 ppm within 20 years and then remains constant at that value. The exact time series of the CO_2_ forcing can be found in Fig. S2.

Plant-FATE does not currently have an explicit representation of nutrient cycles. To simulate different levels of nutrient availability, we assume that plants balance the acquisition of above versus belowground resources (i.e., light and CO_2_ versus nutrients and water) by altering the ratio of fine-root mass to leaf area (or root-to-shoot ratio), *ζ*. While *ζ* can be species-specific in principle, here we use the same value for all species. High *ζ* reflects greater investment in fine roots typical of nutrient-poor habitats, whereas low *ζ* represents fertile soil conditions.

### 7.6 Plant-FATE as a modular vegetation modelling platform

Plant-FATE is not only a new vegetation model, but also a vegetation modelling platform, for the following reasons: (1) Due to its modular design, Plant-FATE can be used to test alternative formulations of core physiological processes. (2) Plant-FATE makes a clean separation between the code for simulating biological processes and the code for simulating population dynamics and trait evolution, thus allowing non-programmers to focus on the biology. To that end, it uses *libpspm*, a numerical library we have developed for solving the partial differential equation underlying the population model in Plant-FATE. (3) Plant-FATE is written in C++ for maximum computational efficiency, with R and Python interfaces providing convenient access for non-specialists. (4) The model is open source, with code available here: https://github.com/jaideep777/Plant-FATE. In future, it will also support massively parallel computing architectures, such as GPUs, making it easier to explore complex ecological questions that require multiple iterations of model runs. Ultimately, Plant-FATE provides a quantitative platform for integrating biodiversity and ecosystem functioning^78^, and thus opens new avenues for understanding the links between biodiversity and ecosystem resilience.

## Supplementary Information captions

### Supplementary text

Plant-FATE model description

### Supplementary figures

Fig. S1. Additional emergent community properties predicted by our model under elevated and ambient CO2 conditions.

Fig. S2. Time series of atmospheric CO2 forcing used for our model runs.

