## Supplementary Text and Figures for "Competition for light can drive adverse species-composition shifts in the Amazon Forest under elevated CO_2_"

**Jaideep Joshi et al.**

July 3, 2023

**A Appendix A: Justification of constant Huber Value within the crown** **27**

**B Supplementary figures** **29**

### 1 Model overview

Plant-FATE predicts the dynamics of multi-species plant communities by scaling the responses of individual plants to the community level. To that end, we (1) describe how individual plants acclimate to their environment by adjusting their physiology (plastic traits) within the bio-physical contraints imposed by their size, architecture, and species-specific (non-plastic) traits; (2) embed individual plants in a physiologically structured population model, where the structured population comprises a diversity of plant species and life-history stages; (3) describe the feedbacks between the population structure and the competitive environment, which govern the outcomes of competition within and among species.

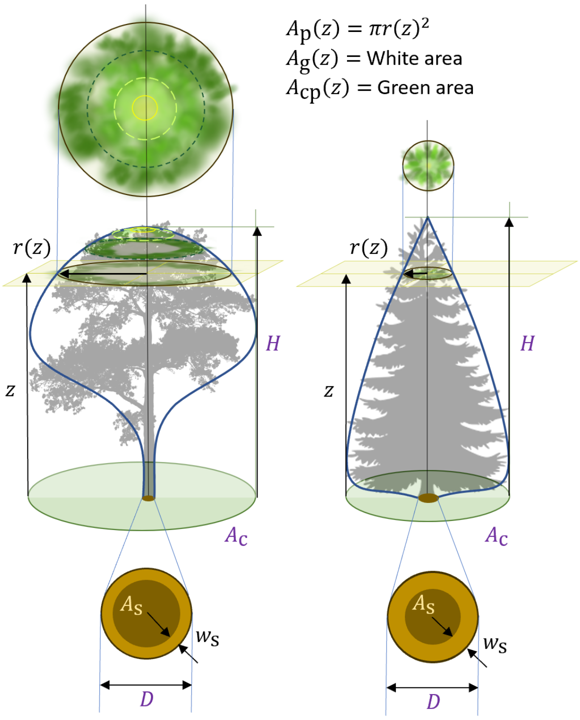

Fig. 1: **Tree architecture**. The basal stem diameter $D$ characterizes tree size in Plant-FATE. It is connected to height $H$, crown projection area $A_c$, basal sapwood area $A_s$, basal sapwood width $w_s$, and other variables via dimensional scaling relationships. Different crown shapes (blue curves) are modelled by specifying the crown radius $r(z)$ at each height $z$. Cross sections of the crown stack together without overlapping to fill the total crown area $A_c$. Each cross-section of the crown contains one or more leaf layers - the number of leaf layers is specified by the crown leaf area index $L$.

### 2 Physiology

In this section, we first describe the architecture of trees in our model. Then, we describe core physiological processes such as photosynthesis, respiration, turnover, and biomass allocation.

#### 2.1 Tree architecture

In this section, we describe the dimensional scaling relationships that connect architectural or size-related variables such as diameter, height, crown area, sapwood area, stem mass, coarse root mass, and fine root mass. Figure 1 summarizes the key concepts used in Plant-FATE for modelling tree architecture. We treat basal diameter as the primary independent variable, connected to all others via scaling relationships. To model these relationships, we extend the T-model[1], as described in further sections.

##### 2.1.1 Height and crown-area scaling (the T-model)

According to the T-model, tree height scales with basal diameter as

$$H = H_m \left(1 - e^{-aD/H_m}\right). \tag{1}$$

If the entire crown is projected onto the ground plane, the area enclosed by the perimeter of the crown (Section 2.1.5, Fig. 1), which we simply refer to as 'crown area' or 'crown projection area', is given by

$$A_c = \frac{\pi c}{4a} HD, \tag{2}$$

where the three parameters $a$, $c$, and $H_m$ are species-specific traits: $H_m$ is the maximum height the tree can attain, $a$ is the 'stem slenderness ratio', i.e., the initial ratio of height to diameter, and $c$ is the ratio of crown projection area to basal sapwood area (see next section).

###### 2.1.2 Sapwood-area and heartwood-area scaling

We define the sapwood fraction $f_s$ as the fraction of the stem cross-sectional area that is sapwood. The heartwood fraction is therefore $f_h = 1 - f_s$. While stem radius may change across the length of the tree, the sapwood fraction is assumed to be constant throughout the tree. However, it may change over time as the tree grows. The ratio of sapwood area to leaf area is called the Huber Value, $\nu_H$, and the ratio of leaf area to crown projection area is the within-crown leaf area index, $L$ (Fig. 1). The ratio of (basal) sapwood area to (projected) crown area is thus

$$\frac{A_s}{A_c} = \frac{A_s}{A_l} \cdot \frac{A_l}{A_c} = \nu_H L. \tag{3}$$

Basal sapwood area is thus given by

$$A_s = \nu_H L A_c = \nu_H L c \left(\frac{\pi}{4} D^2\right) \frac{H}{aD}. \tag{4}$$

The sapwood fraction can then be calculated by dividing sapwood area by the stem cross-sectional area at the tree base,

$$f_s = \frac{A_s}{A_{stem}} = \nu_H L c \frac{H}{aD}. \tag{5}$$

Assuming that sapwood is concentrated within a thin annulus of width $w_s$ at the edge of the stem, the sapwood area is $A_s \approx \pi D w_s$. The sapwood width is therefore,

$$
\begin{aligned}
w_{\mathrm{s}} &= \frac{1}{\pi D}\nu_{\mathrm{H}} L c \left(\frac{\pi}{4} D^2\right) \frac{H}{aD} \\
&= \frac{\nu_{\mathrm{H}} L c}{4a} H_{\mathrm{m}} \left(1 - e^{-aD/H_{\mathrm{m}}}\right) \\
&= \frac{f_{\mathrm{s}}}{4} D
\end{aligned} \tag{6}
$$

Since crown area does not change on short timescales, short-term changes in leaf area are reflected in corresponding changes in Huber value, i.e., $\nu_{\mathrm{H}} L =$ constant in the short term. To determine whether $f_{\mathrm{s}}$ and $\nu_{\mathrm{H}} L$ vary with tree size over longer timescales (i.e., timescales over which the tree grows), we consider two hypotheses for the scaling of sapwood width: (i) $\nu_{\mathrm{H}} L$ is constant throughout the lifetime of the plant, so that sapwood width first increases linearly with diameter and then saturates; or (ii) sapwood fraction is constant throughout the lifetime of the plant, so that sapwood width always increases linearly with diameter. Empirical evidence[2–5] is consistent with the first hypothesis. The T-model also assumes that $\nu_{\mathrm{H}} L c = 1$ throughout the plant lifetime.

To find the value of $\nu_{\mathrm{H}} L$, we note that in young trees, $H = aD$ and $f_{\mathrm{s}} = 1$. Substituting in Eq. 6, we get

$$
\begin{aligned}
\nu_{\mathrm{H}} L &= 1/c \\
f_{\mathrm{s}} &= H/aD \\
f_{\mathrm{h}} &= 1 - H/aD \\
D_{\mathrm{h}} &= \sqrt{4(f_{\mathrm{h}} \pi D^2/4)/\pi} = \sqrt{f_{\mathrm{h}}} D
\end{aligned} \tag{7}
$$

where $D_{\mathrm{h}}$ is heartwood diameter. Equation 7 also implies that heartwood fraction, and consequently heartwood diameter, is approximately zero initially (in the linear region of the $H \sim D$ curve), after which heartwood diameter $D_{\mathrm{h}}$ increases linearly with $D$ (in the saturation region of the $H \sim D$ curve). This is consistent with further empirical observations that show heartwood fraction to be zero until a threshold diameter, followed by a linear increase with diameter[2,3].

##### 2.1.3 Fine root scaling

We assume that total fine root mass $m_{\mathrm{fr}}$ scales *instantaneously* with total leaf area as

$$
m_{\mathrm{fr}} = \zeta A_{\mathrm{l}} = \zeta L A_{\mathrm{c}}, \tag{8}
$$

where $\zeta$ is a species-specific trait.

This instantaneous scaling implies that leaf area and fine-root mass are coordinated. If leaves are shed in response to adverse environmental conditions, fine roots are lost as well, and when new leaves are flushed, there is a corresponding production of fine roots.

##### 2.1.4 Coarse root scaling

Root mass fraction (RMF), i.e., the mass of roots as a fraction of total plant biomass, appears to be a highly conserved quantity across species globally, with median 32% and first and third quantiles at 31% and 34% respectively (Groot database[6]). Therefore, we assume that coarse root mass $m_{\mathrm{cr}}$ is a constant fraction of stem mass $m_{\mathrm{stem}}$,

$$m_{\mathrm{cr}} = f_{\mathrm{cr}} m_{\mathrm{stem}} = f_{\mathrm{cr}} \rho_{\mathrm{s}} V_{\mathrm{s}} / f_{\mathrm{s}}, \tag{9}$$

where $\rho_{\mathrm{s}}$ is the wood density and $V_{\mathrm{s}}$ is sapwood volume. Since leaf and fine-root mass fractions are much smaller than those of stem and coarse roots, $f_{\mathrm{cr}} \approx \frac{RMF}{1-RMF}$. An RMF of 32% implies $f_{\mathrm{cr}} = 0.47$.

##### 2.1.5 Crown shape

Since each leaf must be supplied by water through sapwood, the total volume of sapwood in the tree depends on the distribution of leaves along the tree's height. Most vegetation demographic models assume flat-topped trees, which means that the leaf area is concentrated at the top of the tree. While this simplifies calculations of the vertical light profile, such a flat-topped tree shape is far from reality, and misses important tradeoffs associated with light interception: as we show below, having all leaves at the top would allow trees to capture maximum light, but would require a higher sapwood volume. Therefore, we develop a crown model that allows us to represent more realistic crown shapes and capture the aforementioned trade-offs, while still being tractable enough to yield closed-form expressions for sapwood volume and the vertical light profile.

To that end, we model a radially symmetric crown with cross-sectional radius $r(z)$ at height $z$ (Fig. 1) given by

$$r(z) = r_0 q(z) = r_0 mn \left(\frac{z}{H}\right)^{n-1} \left(1 - \left(\frac{z}{H}\right)^n\right)^{m-1}, \tag{10}$$

where the parameters $m$ and $n$ are species-specific traits. The 3D crown shape is thus a shell obtained by revolving the curve $r(z)$ around the $z$ axis. This function can simulate a wide diversity of crown shapes (Fig. 2). A flat-top model can be recovered by setting $n \to \infty$.

To find the value of $r_0$ in terms of $A_{\mathrm{c}}$, we note that the maximum of $r(z)$ occurs at a height $z_{\mathrm{m}}$ obtained by solving

$$\frac{dr(z)}{dz} = -r_0 Hmn \left(\frac{1}{z^2}\right) \left(\frac{z}{H}\right)^n \left(1 - \left(\frac{z}{H}\right)^n\right)^{m-2} \left((mn-1)\left(\frac{z}{H}\right)^n - (n-1)\right) = 0, \tag{11}$$

which gives

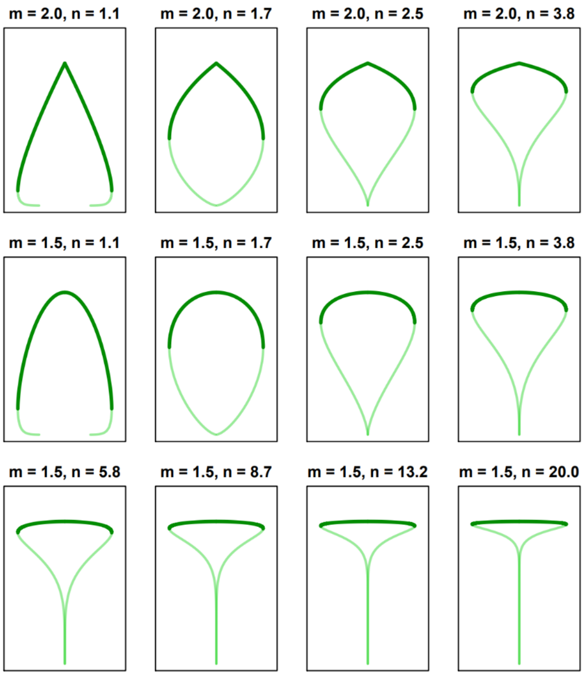

Fig. 2: **Crown shapes as a function of parameters $m$ and $n$.** All trees represented here have the same height and crown area.

$$z_{\mathrm{m}} = H\left(\frac{n-1}{mn-1}\right)^{\frac{1}{n}}. \tag{12}$$

and

$$q_{\mathrm{m}} = q\left(z_{\mathrm{m}}\right) = mn\left(\frac{n-1}{mn-1}\right)^{1-\frac{1}{n}}\left(\frac{(m-1)n}{mn-1}\right)^{m-1}. \tag{13}$$

The maximum crown projection area is thus

$$A_c = \pi r^2\left(z_{\mathrm{m}}\right) = \pi\left(r_0 q\left(z_{\mathrm{m}}\right)\right)^2, \tag{14}$$

giving

$$r_0 = \frac{1}{q_{\mathrm{m}}}\sqrt{\frac{A_c}{\pi}}. \tag{15}$$

To ensure analytical tractability, and to capture the optimality idea that the crown minimizes self-overlap, we make the following additional assumptions about the distribution of leaves through the crown,

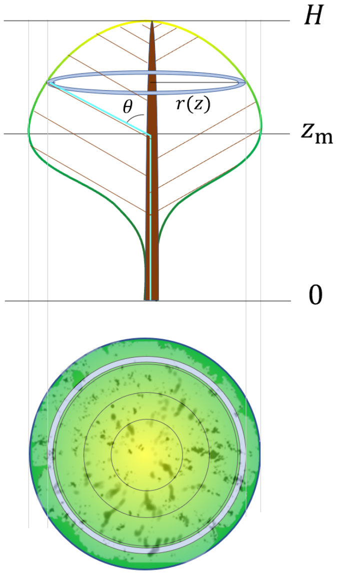

Fig. 3: **Crown-area filling and water supply to leaves**. Figure shows the front view of a vertical section of the crown, together with the top view of the crown. The gradient of colour from dark green to yellow shows the cross sections of the crown at different heights. Most of the leaves lie in the upper crown (height range $[z_{\mathrm{m}}, H]$), with a small area $f_{\mathrm{g}} A_{\mathrm{c}}$ remaining unoccupied (dark specs). This unoccupied area is filled in the lower crown between heights $[0, z_{\mathrm{m}})$ (which is why the specs are dark green and not white). The hydraulic pathway carrying water to leaves located at $(z, r(z))$ first goes up vertically in the trunk, and then through branches at an angle $\theta$ to the trunk.

1. Most leaves are concentrated in the top half of the shell, between $z_{\mathrm{m}}$ and $H$ (the 'upper crown', dark green lines in Fig. 2).

2. A small fraction $f_{\mathrm{g}}$ of the upper crown consists of gaps, such that the upper crown fills up a fraction $1 - f_{\mathrm{g}}$ of the total crown projection area $A_{\mathrm{c}}$.

3. The portion of the crown shell between heights 0 and $z_{\mathrm{m}}$ (the 'lower crown', light green lines in Fig. 2) is mostly empty but contains leaves directly below gaps in the upper crown. Real trees do have a small number of leaves in the lower crown, typically from growing branches. Thus, the projection of the whole crown completely fills the crown projection area $A_{\mathrm{c}}$ (Fig. 3).

4. Leaves in real crowns can experience substantial levels of self-shading. To account for this, we assume that each crown cross-section is composed of $L$ leaf layers. Thus $L$ is the within-crown leaf area index. It is assumed to be constant throughout the crown.

5. Each leaf is supplied by (a network of) xylem vessels through the sapwood. A constant $L$ throughout the crown implies that the sapwood area to leaf area ratio (Huber Value, $\nu_{\mathrm{H}}$) is also constant throughout the crown (but see Appendix A for additional justification).

With these assumptions, we can calculate the sapwood volume. In the derivations below, we start without the assumption of constant $L$ and $\nu_{\mathrm{H}}$ – we treat them as functions of $z$, and set them to constants only when needed.

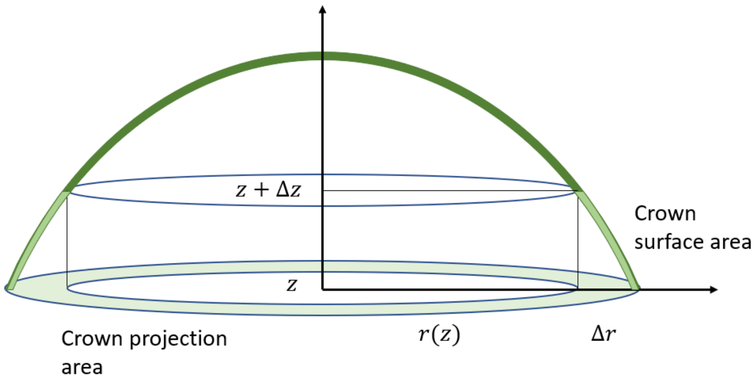

Fig. 4: Crown section

Consider a cross-section of the upper crown between heights $z$ and $z + \Delta z$. The projected crown from this cross-section is distributed (potentially with gaps) in an annulus of radius $r(z)$ and width $\Delta r$ (Fig. 4). The filled projection area of the crown within this layer therefore is

$$\begin{aligned} \Delta A_{\mathrm{cp}}(z) &= \pi \left( r(z)^2 - r(z+\Delta z)^2 \right) (1 - f_{\mathrm{g}}) \\ &\approx 2\pi r(z) \cdot \Delta r \cdot (1 - f_{\mathrm{g}}) \\ &= 2\pi r(z) \cdot (1 - f_{\mathrm{g}}) \cdot \left| \frac{dr}{dz} \right| \Delta z = 2\pi \left(1 - f_{\mathrm{g}}\right) r(z) \left| r'(z) \right| \Delta z. \end{aligned} \tag{16}$$

If the leaf area index in this cross section is $L(z)$, then the leaf area contained within it is

$$\Delta A_{\mathrm{l}}(z) = L(z) \Delta A_{\mathrm{cp}}(z) = 2\pi \left(1 - f_{\mathrm{g}}\right) L(z) r(z) \left| r'(z) \right| \Delta z, \tag{17}$$

and the area of sapwood needed to supply water to the leaves in this layer is

$$\Delta A_{\mathrm{s}} = \nu_{\mathrm{H}}(z) L(z) \Delta A_{\mathrm{cp}}(z) = 2\pi \left(1 - f_{\mathrm{g}}\right) \nu_{\mathrm{H}}(z) L(z) r(z) \left| r'(z) \right| \Delta z. \tag{18}$$

If the length of the hydraulic pathway (consisting of the network of xylem vessels and surrounding tissue) supplying water to the leaves in this layer is $l(z)$, its volume is

$$\Delta V_{\mathrm{s}} = l(z) \Delta A_{\mathrm{s}}(z) = 2\pi \left(1 - f_{\mathrm{g}}\right) \nu_{\mathrm{H}}(z) L(z) r(z) \left| r'(z) \right| l(z) \Delta z \tag{19}$$

The total volume of sapwood in the tree is obtained by integrating Eq. 19 over the upper and lower canopy,

$$
\begin{aligned}
V_{\mathrm{s}} &= V_{\mathrm{sl}} + V_{\mathrm{su}} \\
&= \int_0^{z_{\mathrm{m}}} 2\pi f_{\mathrm{g}} \nu_{\mathrm{H}}(z) L(z) r(z) r'(z) l(z) dz - \int_{z_{\mathrm{m}}}^{H} 2\pi \left(1 - f_{\mathrm{g}}\right) \nu_{\mathrm{H}}(z) L(z) r(z) r'(z) l(z) dz,
\end{aligned} \tag{20}
$$

Note that the second (upper canopy) term has a negative sign because $r'(z)$ is negative in the upper canopy.

To solve for $V_{\mathrm{s}}$, we first evaluate the indefinite integral

$$
I(z) = \int \nu_{\mathrm{H}}(z) L(z) r(z) r'(z) l(z) dz. \tag{21}
$$

In principle, the variations in $L(z)$ over height could result from different light levels experienced by leaves at different heights. However, explicitly keeping track of the vertical distribution of $L(z)$ is extremely computationally expensive. Therefore, we assume a constant average $L$ throughout the crown. This also implies a constant average $\nu_{\mathrm{H}}(z)$. We further assume that the vessels carrying water to leaves located at $(r(z), z)$ first go up vertically within the trunk and then travel through branches that are at an angle $\theta$ to the trunk (cyan line in Fig. 3). To calculate the total path length $l(z)$, note that the path length within the trunk is

$$
l_{\mathrm{trunk}}(z) = z - \frac{r(z)}{\tan(\theta)}, \tag{22}
$$

and the path length within branches is

$$
l_{\mathrm{branch}}(z) = \frac{r(z)}{\sin(\theta)}. \tag{23}
$$

Thus, the total path length is

$$
\begin{aligned}
l(z) &= z + r(z) \left( \frac{(1 - \cos(\theta))}{\sin(\theta)} \right) \\
&= z + r(z) \left( \frac{2 \sin^2(\theta)}{2 \cos(\theta/2) \sin(\theta/2)} \right) \\
&= z + r(z) \tan\left(\theta/2\right).
\end{aligned} \tag{24}
$$

Substituting in Eq. 21, we get

$$I = L\nu_{\mathrm{H}} \int \left( z + r(z) \tan\left(\frac{\theta}{2}\right)\right) r(z) r'(z) dz. \tag{25}$$

Using the identities $dr = r'(z)dz$ and $\frac{1}{2}d\left(r^2(z)\right) = r(z)r'(z)dz$, we get

$$I = L\nu_{\mathrm{H}} \left( \frac{1}{2} \int zd(r^2) \ + \tan\left(\frac{\theta}{2}\right) \int r^2(z) dr \right). \tag{26}$$

Integrating the first term by parts ($\int udv = uv - \int vdu$, with $u = z$ and $v = r^2(z)$), we get

$$I(z) = \frac{1}{2} L\nu_{\mathrm{H}} \ \left( zr^2(z) - \int r^2(z)dz + 2\tan\left(\frac{\theta}{2}\right) \frac{r^3(z)}{3}\right). \tag{27}$$

Further,

$$\int r^2(z)dz = r_0^2 \int q^2(z)dz = Hr_0^2 m^2 n \cdot B\left( \left(\frac{z}{H}\right)^n ; 2 - \frac{1}{n},\ 2m-1 \right), \tag{28}$$

where $B(x; p, q)$ is the un-normalized incomplete Beta function evaluated at $x$.

Now, let $\alpha(z) = zr^2(z)$, $\beta(z) = \int r^2(z)dz$, and $\gamma(z) = 2\tan\left(\frac{\theta}{2}\right) \frac{r^3(z)}{3}$. Then,

$$I(z) = \frac{1}{2} L\nu_{\mathrm{H}} \left(\alpha(z) - \beta(z) + \gamma(z)\right),$$

with $\alpha(0) = \alpha(H) = 0$, $\beta(0) = 0$, and $\gamma(0) = \gamma(H) = 0$.

Substituting in Eq. 20, we get

$$
\begin{aligned}
V_{\mathrm{s}} &= 2\pi \left(f_{\mathrm{g}} \left(I\left(z_{\mathrm{m}}\right) - I(0)\right) - \left(1 - f_{\mathrm{g}}\right)\left(I(H) - I\left(z_{\mathrm{m}}\right)\right)\right) \\
&= \pi L \nu_{\mathrm{H}} \left(\alpha\left(z_{\mathrm{m}}\right) - \beta\left(z_{\mathrm{m}}\right) + \left(1 - f_{\mathrm{g}}\right)\beta(H) + \gamma\left(z_{\mathrm{m}}\right)\right) \\
&= \pi L \nu_{\mathrm{H}} \Bigg( z_{\mathrm{m}} r_0^2 q_{\mathrm{m}}^2 - H r_0^2 m^2 n \cdot B\left(\left(\frac{z_{\mathrm{m}}}{H}\right)^n; 2 - \frac{1}{n},\ 2m - 1\right) \\
&\qquad + \left(1 - f_{\mathrm{g}}\right) H r_0^2 m^2 n \cdot B\left(1; 2 - \frac{1}{n},\ 2m - 1\right) \\
&\qquad + 2\tan\left(\frac{\theta}{2}\right) \frac{r_0^3 q_{\mathrm{m}}^3}{3} \Bigg) \\
&= L \nu_{\mathrm{H}} H A_{\mathrm{c}} \Bigg( \left(\frac{n-1}{mn-1}\right)^{1/n} \\
&\qquad - \frac{m^2 n}{q_{\mathrm{m}}^2} \cdot B\left(\frac{n-1}{mn-1}; 2 - \frac{1}{n},\ 2m - 1\right) \\
&\qquad + \left(1 - f_{\mathrm{g}}\right) \frac{m^2 n}{q_{\mathrm{m}}^2} \cdot B\left(1; 2 - \frac{1}{n}, 2m - 1\right) \\
&\qquad + 2\tan\left(\frac{\theta}{2}\right) \frac{A_{\mathrm{c}}^{1/2}}{3\sqrt{\pi} H} \Bigg)
\end{aligned}
\tag{29}
$$

Note that the first three terms are dependent on only $m$ and $n$, and can thus be precomputed (Fig. 5). Denoting these by $\eta_c(m, n)$, and assuming the tilt angle of branches to be 90°, we get

$$
\begin{aligned}
V_{\mathrm{s}} =&\ L \nu_{\mathrm{H}} H A_{\mathrm{c}} \left(\eta_c(m, n) + \frac{2}{3H}\sqrt{\frac{A_{\mathrm{c}}}{\pi}}\right) \\
&= \frac{H A_{\mathrm{c}}}{c} \left(\eta_c(m, n) + \frac{2}{3H}\sqrt{\frac{A_{\mathrm{c}}}{\pi}}\right) \\
&= A_{\mathrm{s}} H \left(\eta_c(m, n) + \frac{2}{3H} r_{\mathrm{c}}\right).
\end{aligned}
\tag{30}
$$

The term $\eta_{\mathrm{c}}$ defines the taper in the total cross-sectional area (trunk + branches) with height. For e.g., for flat-topped trees, $\eta_{\mathrm{c}} = 1$, which means that the stem is cylindrical, and has the highest volume. For bottom-heavy trees like conifers, $\eta_{\mathrm{c}} \approx 0.5$, which implies a conical or paraboloid stem. This elucidates the aforementioned tradeoffs between light interception (by placing leaves as high as possible) and avoiding excessive sapwood costs (by placing the leaves lower and thus reducing sapwood volume).

#### 2.2 Photosynthesis, respiration, and turnover

Gross $CO_2$ assimilation rate depends on the intensity of incident light $I_0$ in the direction perpendicular to the crown, the fraction $f_{\mathrm{apar}}$ of this incident light absorbed by the leaves in that cross-section, and other environmental variables such as atmospheric $CO_2$ concentration, vapour pressure deficit (VPD), temperature, and atmospheric pressure.

We calculate photosynthetic traits and photosynthesis rate per unit crown area by approximating each crown cross-section with a 'big leaf', i.e., the fraction of light absorbed collectively by leaves

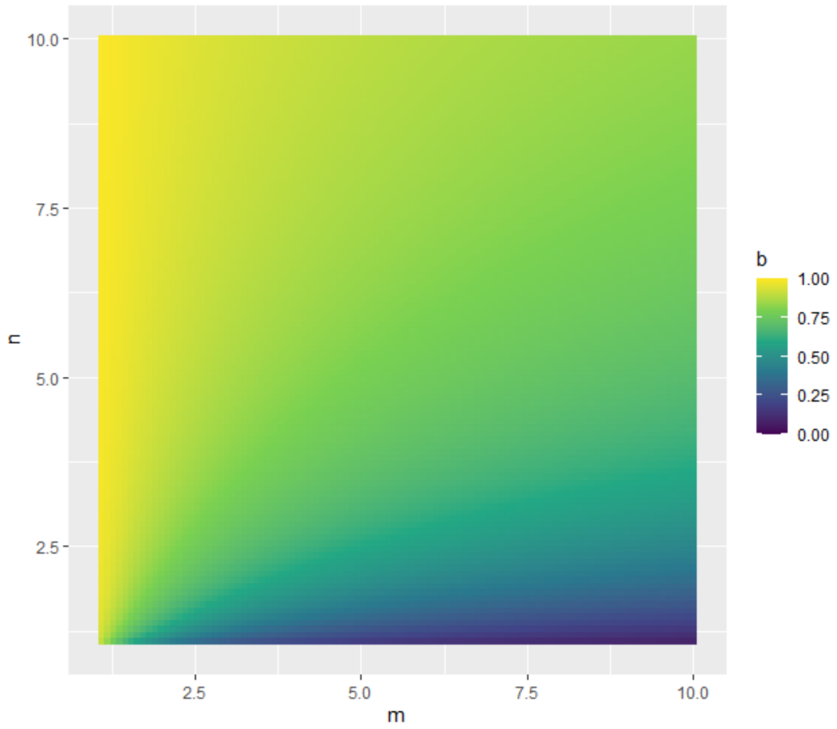

Fig. 5: $\eta_c(m, n)$

in each cross-section is

$$f_{\text{apar}} = 1 - e^{-kL}, \tag{31}$$

where $k$ is the light extinction coefficient.

To calculate the crown-area-specific photosynthetic capacities ($V_{\text{cmax}}$, $J_{\text{max}}$), stomatal conductance $g_{\text{c}}$, and photosynthesis rate $P_{\text{c}}$, we use P-hydro[7], a trait-based model that uses optimality principles to predict the simultaneous acclimation of photosynthetic capacity and stomatal conductance on daily and weekly timescales. However, in this work, we run the model only on weekly (and above) timescales, as follows.

First, the acclimated photosynthetic capacities and maximum gas exchange rates ($g_{\text{c,max}}$ and $P_{\text{c,max}}$) are calculated based on the monthly maximum light intensity, $I_{\text{max}}$. Then, the monthly average value of photosynthesis rate is calculated as

$$P_{\text{c}} = 1.18 P_{\text{c,max}} \frac{I_{\text{avg}}}{I_{\text{max}}}, \tag{32}$$

where the factor 1.18 accounts for the slight nonlinearity in the light response of instantaneous photosynthesis rates, and has been calculated using the light response predicted by the P-hydro model.

The gross whole-plant $CO_2$ update rate is thus

$$P_{\text{gross}} = P_{\text{c}} A_{\text{c}}. \tag{33}$$

Net whole-plant biomass production rate is calculated as

$$P_{\text{net}} = c_{\text{bio}} y \left(P_{\text{gross}} - R\right) - T \tag{34}$$

where $c_{\text{bio}}$ is the biomass produced per mol $CO_2$, $y$ is the yield factor accounting for biomass lost in growth respiration, $R$ is the maintenance respiration rate of leaves, fine roots, and wood, and $T$ is the turnover rate of leaves and fine roots.

The total plant respiration rate is the sum of the respiration rates $R_{\text{s}}$, $R_{\text{r}}$, and $R_{\text{l}}$ of sapwood, fine roots, and leaves, respectively. Following empirical observations, sapwood respiration rate is assumed to scale with sapwood mass[8], fine root respiration rate is assumed to scale with fine root mass as well as gross productivity[9–11], and leaf respiration rate is assumed to be proportional to the photosynthetic capacity $V_{\text{cmax}}$[12]:

$$\begin{aligned} R_{\text{s}} &= r_{\text{s}} m_{\text{s}} \\ R_{\text{r}} &= r_{\text{r}} m_{\text{fr}} P_{\text{c}} \\ R_{\text{l}} &= r_{\text{dark}} V_{\text{cmax}} A_{\text{c}}, \\ R &= R_{\text{s}} + R_{\text{r}} + R_{\text{l}}. \end{aligned} \tag{35}$$

The sapwood mass required in the above equation is given by,

$$\begin{aligned} m_{\text{s}} &= \rho_{\text{s}} A_{\text{s}} H \left(\eta_c(m,n) + \frac{2}{3H} r_{\text{c}}\right) \\ &= \rho_{\text{s}} \frac{A_{\text{c}}}{c} H \left(\eta_c(m,n) + \frac{2}{3H}\sqrt{\frac{A_{\text{c}}}{\pi}}\right). \end{aligned} \tag{36}$$

Turnover rates of leaves and fine roots are calculated from the masses ($m_{\text{l}}$ and $m_{\text{fr}}$, respectively) and lifespans ($\tau_{\text{l}}$ and $\tau_{\text{fr}}$, respectively) of these pools,

$$T = \frac{m_{\text{fr}}}{\tau_{\text{fr}}} + \frac{m_{\text{l}}}{\tau_{\text{l}}}, \tag{37}$$

#### 2.3 Optimal leaf and fine-root economics

To calculate the leaf and fine-root lifespans ($\tau_{\text{l}}$ and $\tau_{\text{fr}}$, respectively), we adopt the model of optimal leaf economics (LES) for evergreen species developed in ref[13], which predicts optimal leaf lifespan in a given (fixed) environment. The same calculations also hold for ergodic environments, as long as the variance in environmental variables is small. Furthermore, since we are primarily concerned with a largely aseasonal site in the Amazon rainforest in this work, we also neglect seasonality and assume that the growing season spans the entire year.

The central idea for modelling leaf lifespan is that the leaf starts with a certain photosynthetic capacity which decreases with leaf age, and the leaf is dropped when returns on investment are no longer sustainable. The optimal-LES theory assumes that crown-area-specific photosynthesis rate decreases with leaf age as $P_c(t) = P_{c,0}(1 - t/b)$. Thus, for evergreen leaves, the average photosynthesis rate per unit crown area over the entire leaf lifecycle is

$$P_{c,avg} = \frac{1}{\tau_l}\int_0^{\tau_l} P_{c,0}(1 - t/b)dt = P_{c,0}\left(1 - \frac{\tau_l}{2b}\right) \tag{38}$$

where $P_{c,0} = k_1 k_2 V_{cmax} m_c$ is the instantaneous photosynthesis rate per unit crown area ($k_1$ and $k_2$ convert the units of $V_{cmax}$ from $\mu\text{mol} \cdot \text{m}^{-2} \cdot \text{s}^{-1}$ to $\text{g} \cdot \text{m}^{-2} \cdot \text{day}^{-1}$). Here,

$$m_c = \frac{c_i - \Gamma^*}{c_i + K_M} \tag{39}$$

where $c_i$ is the leaf internal $CO_2$ concentration, $\Gamma^*$ is the light compensation point, and $K_M$ is the Michaelis-Menten coefficient.

The net profit for the plant (per unit crown area) is the average photosynthesis rate minus the amortized construction and maintenance costs of leaves and supporting tissues (including fine roots; per unit crown area),

$$\text{Profit} = P_{c,avg} - \frac{C_c \lambda_l L}{\tau_l}, \tag{40}$$

where $C_c$ represents the costs per unit leaf mass and $\lambda_l$ is the leaf mass per unit leaf area (LMA), which is a species-specific trait.

The leaf lifespan that maximises profit is calculated by setting the derivative of profit with respect to $\tau_l$ to zero,

$$\begin{aligned} \frac{d\text{Profit}}{d\tau_l} = 0 &\implies -\frac{P_{c,0}}{2b} + \frac{C_c L \lambda_l}{\tau_l^2} = 0 \\ &\implies \tau_l^* = \sqrt{\frac{2bC_c L\lambda_l}{P_{c,0}}} \end{aligned} \tag{41}$$

Further, it has been empirically observed[14] that

$$b = \frac{u\lambda_{\mathrm{l}}}{k_1 k_2 V_{\mathrm{cmax,25,l}}} \tag{42}$$

where $V_{\mathrm{cmax,25,l}}$ stands for photosynthetic capacity per unit leaf area at $25^o$C. To express it in terms of $V_{\mathrm{cmax,25}}$ (i.e., photosynthetic capacity per unit crown area at $25^o$C), we note that $V_{\mathrm{cmax,l}} = V_{\mathrm{cmax}} A_{\mathrm{c}}/A_{\mathrm{l}} = V_{\mathrm{cmax}}/L$. Thus,

$$b = \frac{u\lambda_{\mathrm{l}} L}{k_1 k_2 V_{\mathrm{cmax,25}}} \tag{43}$$

and thus, optimal leaf lifespan is

$$\begin{aligned}
\tau_{\mathrm{l}}^* &= \sqrt{\frac{2uC_{\mathrm{c}}L^2\lambda_{\mathrm{l}}^2}{k_1k_2V_{\mathrm{cmax,25}}P_{\mathrm{c,0}}}} \\
&= \sqrt{\frac{2uC_{\mathrm{c}}h_{\mathrm{T}}L^2\lambda_{\mathrm{l}}^2}{k_1k_2V_{\mathrm{cmax}}k_1k_2V_{\mathrm{cmax}}m_{\mathrm{c}}}} \\
&= \frac{\lambda_{\mathrm{l}}L}{V_{\mathrm{cmax}}}\sqrt{\frac{2uC_{\mathrm{c}}h_{\mathrm{T}}}{(k_1k_2)^2m_{\mathrm{c}}}} = \frac{\lambda_{\mathrm{l}}L}{V_{\mathrm{cmax,25}}}\sqrt{\frac{2uC_{\mathrm{c}}}{(k_1k_2)^2m_{\mathrm{c}}h_{\mathrm{T}}}}. \\
&= \frac{\lambda_{\mathrm{l}}L}{V_{\mathrm{cmax,25}}} f_{\mathrm{LES}}(\chi, T)
\end{aligned} \tag{44}$$

where the factor $h_{\mathrm{T}}$ is the arrhenius temperature response of $V_{\mathrm{cmax}}$ (i.e., $V_{\mathrm{cmax}} = V_{\mathrm{cmax,25}}h_{\mathrm{T}}$). The factor $f_{\mathrm{LES}}(\chi, T)$, which is a function of temperature $T$ and the leaf internal-to-external $CO_2$ ratio $\chi$, captures the environmental response of leaf lifespan,

$$f_{\mathrm{LES}}(\chi, T) = \sqrt{\frac{2uC_{\mathrm{c}}}{(k_1k_2)^2m_{\mathrm{c}}h_{\mathrm{T}}}} \tag{45}$$

The optimal average photosynthesis rate is thus

$$\begin{aligned}
P_{\mathrm{c,avg}} &= P_{\mathrm{c,0}}\left(1 - \frac{\tau_{\mathrm{l}}}{2b}\right) \\
&= P_{\mathrm{c,0}}\left(1 - \frac{\frac{\lambda_{\mathrm{l}}L}{V_{\mathrm{cmax}}}\sqrt{\frac{2uC_{\mathrm{c}}h_{\mathrm{T}}}{(k_1k_2)^2m_{\mathrm{c}}}}}{2\frac{u\lambda_{\mathrm{l}}Lh_{\mathrm{T}}}{k_1k_2V_{\mathrm{cmax}}}}\right) \\
&= P_{\mathrm{c,0}}\left(1 - \frac{\sqrt{\frac{2uC_{\mathrm{c}}h_{\mathrm{T}}}{(k_1k_2)^2m_{\mathrm{c}}}}}{\frac{2uh_{\mathrm{T}}}{k_1k_2}}\right) \\
&= P_{\mathrm{c,0}}\left(1 - \sqrt{\frac{C_{\mathrm{c}}}{2uh_{\mathrm{T}}m_{\mathrm{c}}}}\right).
\end{aligned} \tag{46}$$

This means, a correction factor of $1-\sqrt{\frac{C_\mathrm{c}}{2uh_\mathrm{T}m_\mathrm{c}}}$ must be applied to the photosynthesis rate calculated for newborn leaves, to account for average leaf age.

The cost of construction and maintenance of leaves and supporting tissues (i.e., fine roots, stem, etc) is

$$\begin{aligned}C_\mathrm{leaf} &= C_\mathrm{c}\lambda_\mathrm{l}L/\tau_\mathrm{l} \\ &= C_\mathrm{c}V_\mathrm{cmax}k_1k_2\sqrt{\frac{m_\mathrm{c}}{2uC_\mathrm{c}h_\mathrm{T}}} \\ &= V_\mathrm{cmax}k_1k_2m_\mathrm{c}\sqrt{\frac{C_\mathrm{c}}{2um_\mathrm{c}h_\mathrm{T}}} \\ &= V_\mathrm{cmax}\sqrt{\frac{(k_1k_2)^2C_\mathrm{c}m_\mathrm{c}}{2uh_\mathrm{T}}} \\ &= P_\mathrm{c,0}\sqrt{\frac{C_\mathrm{c}}{2um_\mathrm{c}h_\mathrm{T}}}\end{aligned} \tag{47}$$

Thus, net profit is

$$\mathrm{Profit} = P_\mathrm{c,0}\left(1-\sqrt{\frac{2C_\mathrm{c}}{uh_\mathrm{T}m_\mathrm{c}}}\right) \tag{48}$$

The P-hydro model of photosynthesis used in Plant-FATE (Section 2.2) is calibrated with assimilation rates measured on average leaves (not young leaves), and thus the photosynthesis rate $P_\mathrm{c}$ obtained from it corresponds to the average photosynthesis rate $P_\mathrm{c,avg}$ considered here,

$$P_\mathrm{c,avg} = P_\mathrm{c} \tag{49}$$

Furthermore, we do not subtract the cost of leaf construction and maintenance (Eq. 47) from $P_\mathrm{c,avg}$ because carbon allocated to construction and carbon lost by maintenance respiration is accounted for explicitly (Section 2.2).

Empirical studies on root economics suggest that fine-root lifespan is negatively correlated with specific root length (SRL or $\sigma_\mathrm{r}$). Thus, SRL is the analogue of SLA for roots. Without formulating an explicit optimality model for fine roots, we hypothesise here that leaves and fine roots respond to the environment in a coordinated way. Therefore, we model fine-root lifespan analogously to Eq. 44,

$$\tau_\mathrm{r}^* = c_\mathrm{lr}\frac{(1/\sigma_\mathrm{r})}{V_\mathrm{cmax,25}}f_\mathrm{LES}(\chi, T) \tag{50}$$

where $c_\mathrm{lr}$ is a constant.

#### 2.4 Biomass Allocation

The architectural constraints and dimensional relationships derived above lead to a model of biomass partitioning among different organs. The total rate of biomass production, to be allocated to various carbon pools, is

$$\frac{dB}{dt} = \max\left(P_{\text{net}}, 0\right), \tag{51}$$

where $P_{\text{net}}$ is the net whole-plant photosynthesis rate. If $P_{\text{net}}$ is negative, there is no biomass production.

To calculate the change in a dimensional variable $X$, we may need to use the chain rule,

$$\frac{dX}{dt} = \frac{dX}{dD}\frac{dD}{dt}. \tag{52}$$

Therefore we first calculate the derivatives of each dimensional variable with respect to diameter. The derivatives of height and crown area with respect to diameter are

$$\begin{aligned} \frac{dH}{dD} &= ae^{-\frac{aD}{H_{\text{m}}}} \\ \frac{dA_{\text{c}}}{dD} &= \frac{\pi c}{4a}\left(D\frac{dH}{dD} + H\right). \end{aligned} \tag{53}$$

To find the mass of stem and branches, we note that

$$\begin{aligned} V_{\text{s}} &= A_{\text{s,base}} H \left(\eta_c(m,n) + \frac{2}{3H} r_{\text{c}}\right) \\ &= \ H \cdot \frac{\pi D^2}{4} \cdot f_{\text{s}} \left(\eta_c(m,n) + \frac{2}{3H} r_{\text{c}}\right) \end{aligned} \tag{54}$$

$$\begin{aligned} m_{\text{stem}} &= \rho_{\text{s}} H \cdot \frac{\pi D^2}{4} \cdot \left(\eta_c(m,n) + \frac{2}{3H} r_{\text{c}}\right) \\ &= \rho_{\text{s}} \frac{\pi D^2 H}{4} \eta_c(m,n) + \rho_{\text{s}} \frac{\pi D^2}{6} \sqrt{\frac{c}{4a} DH} \\ &= \rho_{\text{s}} \frac{\pi D^2 H}{4} \eta_c(m,n) + \rho_{\text{s}} \frac{\pi D^2}{6} \sqrt{\frac{c}{4a} DH} \\ &= \rho_{\text{s}} \frac{\pi}{4} \eta_c(m,n) D^2 H + \rho_{\text{s}} \frac{\pi}{12} \sqrt{\frac{c}{a}} D^{5/2} H^{1/2}. \end{aligned} \tag{55}$$

The first term on the RHS represents the mass of the trunk $m_{\text{trunk}}$, and the second term represents the mass of branches $m_{\text{branch}}$. The derivatives of trunk and branch masses with diameter are then

$$
\begin{aligned}
\frac{dm_{\text{trunk}}}{dD} &= \rho_{\text{s}} \frac{\pi}{4} \eta_c(m,n) \left( D^2 \frac{dH}{dD} + 2DH \right), \\
\frac{dm_{\text{branch}}}{dD} &= \rho_{\text{s}} \frac{\pi}{12} \sqrt{\frac{c}{a}} \left( \frac{1}{2} D^{5/2} H^{-1/2} \frac{dH}{dD} + \frac{5}{2} D^{3/2} H^{1/2} \right).
\end{aligned}
\tag{56}
$$

The derivative of coarse root mass with respect to diameter is

$$
\frac{dm_{\text{cr}}}{dD} = f_{\text{cr}} \left( \frac{dm_{\text{trunk}}}{dD} + \frac{dm_{\text{branch}}}{dD} \right). \tag{57}
$$

The rate of change of leaf mass $m_{\text{l}}$ is

$$
\begin{aligned}
\frac{dm_{\text{l}}}{dt} &= \frac{d}{dt} \left( \lambda_{\text{l}} L A_{\text{c}} \right) \\
&= \lambda_{\text{l}} L \frac{dA_{\text{c}}}{dD} \frac{dD}{dt} + \lambda_{\text{l}} A_{\text{c}} \frac{dL}{dt} \\
&= \frac{dm_{\text{l,growth}}}{dD} \frac{dD}{dt} + \frac{dm_{\text{l,lai}}}{dt},
\end{aligned}
\tag{58}
$$

where the first term on the RHS represents the increment in leaf mass due to plant growth, and the second term represents the changes in leaf mass due to changes in crown LAI.

The rate of change of fine-root mass $m_{\text{fr}}$ is

$$
\frac{dm_{\text{fr}}}{dt} = \frac{d}{dt} \left( \frac{\zeta}{\lambda_{\text{l}}} m_{\text{l}} \right) = \frac{\zeta}{\lambda_{\text{l}}} \frac{dm_{\text{l}}}{dt}. \tag{59}
$$

Note that this implies that a change in LAI instantaneously leads to a coordinated change in change in fine-root mass. This is not incompatible with fact that leaf and fine-root lifespans are unequal – it just means that at any given point in time, leaf and fine-root masses maintain the ratio $\lambda_{\text{l}}/\zeta$, even though their turnover rates may be different.

Since LAI changes are fundamental to future photosynthesis and preventing mortality, we first subtract the biomass allocated to LAI change from total production. The remaining biomass is partitioned into reproduction and growth. For a given rate of change of LAI ($dL/dt$), the biomass required to bring about this LAI change is given by

$$
\frac{dm_{\text{lai}}}{dt}|_{\text{req}} = (\lambda_{\text{l}} + \zeta) \cdot A_{\text{c}} \cdot \frac{dL}{dt}. \tag{60}
$$

However, whereas an increase in LAI requires additional biomass investment into leaves and fine roots, a decrease in LAI simply results from leaf and fine-root shedding. Furthermore, the biomass allocated to LAI change cannot exceed (a fraction $f_{\text{L}}$ of) total production. Therefore, the actual biomass allocation to LAI change is

$$\frac{dm_{\text{lai}}}{dt} = \begin{cases} \min\left((\lambda_l + \zeta)\cdot A_c \cdot \frac{dL}{dt},\ \ f_L \frac{dB}{dt}\right), & \frac{dL}{dt} > 0 \\ 0, & \frac{dL}{dt} \le 0 \end{cases} \tag{61}$$

If the required allocation exceeds $f_L \frac{dB}{dt}$, $dL/dt$ is recalculated as

$$\frac{dL}{dt}|_{\max} = \frac{f_L}{(\lambda_l + \zeta)\cdot A_c}\frac{dB}{dt}. \tag{62}$$

The remaining biomass is allocated for growth and reproduction,

$$\frac{dm_{\text{gr}}}{dt} = \frac{dB}{dt} - \frac{dm_{\text{lai}}}{dt}, \tag{63}$$

out of which a fraction $f_r(D)$ is allocated to reproduction. Thus, the biomass allocated for growth is

$$\frac{dm_g}{dt} = (1 - f_r(D))\,\frac{dm_{\text{gr}}}{dt}. \tag{64}$$

The fraction of biomass allocated to reproduction is given by

$$f_r(D) = f_{r,\max}\frac{1}{1 + e^{a_2(1 - D/D_{\text{mat}})}}, \tag{65}$$

where $f_{r,\max}$ is the maximum fraction of biomass allocated to reproduction, $a_2$ is the rate at which allocation to reproduction increases as the tree grows and matures, and $D_{\text{mat}}$ is the diameter at which reproductive allocation reaches 50% of its maximum. This diameter $D_{\text{mat}}$ is taken to be the diameter when the tree height reaches $f_{\text{hmat}} H_m$, where $f_{\text{hmat}}$ is a constant fraction.

The biomass allocated to growth leads to an increment in diameter, as well as increments in the masses of leaves, fine roots, trunk, branches, and coarse roots,

$$\frac{dm_g}{dt} = \left(\frac{dm_{\text{l,growth}}}{dD}\left(1 + \frac{\zeta}{\lambda_l}\right) + \left(\frac{dm_{\text{trunk}}}{dD} + \frac{dm_{\text{branch}}}{dD}\right)(1 + f_{\text{cr}})\right)\frac{dD}{dt}. \tag{66}$$

### 3 Community dynamics

The allocation model described above allows us to describe the demographic rates of plants (size growth rate, mortality rate, and fecundity rate) as functions of plant size $s$, traits $x$, time $t$, and the environment $E$. These demographic rates define the physiologically structured population model, which forms the core of Plant-FATE.

#### 3.1 Growth, mortality, and fecundity rates

In Plant-FATE, size is defined by basal diameter (therefore, $s := D$). Size growth rate $g(s,x,t,E) = ds/dt$ can be obtained by simply rearranging Eq. 66,

$$g(s,x,t,E) = \left( \frac{dm_{\mathrm{l,growth}}}{dD} \left( 1 + \frac{\zeta}{\lambda_{\mathrm{l}}} \right) + \left( \frac{dm_{\mathrm{trunk}}}{dD} + \frac{dm_{\mathrm{branch}}}{dD} \right) (1 + f_{\mathrm{cr}}) \right)^{-1} \frac{dm_{\mathrm{g}}}{dt}. \tag{67}$$

Fecundity rate $f(s,t,E)$, defined as the number of seeds produced per unit time, can be calculated from the biomass allocated to reproduction,

$$f(s,x,t,E) = \frac{1}{m_{\mathrm{seed}}(1 + c_{\mathrm{acc}})} \left( f_{\mathrm{r}}(s) \frac{dm_{\mathrm{gr}}}{dt} \right), \tag{68}$$

where $m_{\mathrm{seed}}$ is the seed mass and $c_{\mathrm{acc}}$ is the mass of seed accessories per unit seed mass.

We use an empirically motivated mortality rate function $\mu(s,x,t,E)$, which accounts for the decrease in mortality with wood density and productivity (the 'growth-mortality tradeoff'), an initial decrease in mortality with diameter which captures improvements in seedling survival with size, and a subsequent increase in mortality with increasing diameter, which captures the accelerated death of very large trees.

$$\mu(s,x,t,E) = \gamma_{\mathrm{m}} \left( \frac{\rho_{\mathrm{s}}}{\rho_{\mathrm{s,0}}} \right)^{\epsilon_\gamma} + \alpha_{\mathrm{m}} \left( \frac{\rho_{\mathrm{s}}}{\rho_{\mathrm{s,0}}} \right)^{\epsilon_\alpha} e^{-\beta_{\mathrm{m}} g(s,x,t,E)} + c_{\mathrm{D,0}} \left( \frac{s}{D_0} \right)^{\epsilon_{\mathrm{d0}}} \left( \frac{\rho_{\mathrm{s}}}{\rho_{\mathrm{s,0}}} \right)^{\epsilon_{\mathrm{D}}} + c_{\mathrm{D,1}} e^{-s/D_{\mathrm{a}}} \tag{69}$$

#### 3.2 The physiologically structured population model

The population model describes the time-evolution of the size distribution of individuals in the population. It is physiologically structured because the demographic rates depend on size and traits – thus, individuals of different species and developmental stages differ in their demographic rates. The dynamics of the population is described by the McKendrick-von Foerster equation,

$$\frac{\partial u(s,x,t)}{\partial t} = \frac{\partial}{\partial s} \left( g(s,x,t,E) u(s,x,t) \right) - \mu(s,x,t,E) u(s,x,t), \tag{70}$$

where $u(s,x,t)$ is the density of individuals with size $s$ and traits $x$ at time $t$, i.e., the size interval $[s,\ s+ds)$ contains $u(s,x,t)ds$ individuals with traits $x$ per unit ground area.

The partial differential equation (PDE; Eq. 70) has the following boundary condition,

$$g\left(s_{\mathrm{b}}, x, t, E\right) u\left(s_{\mathrm{b}}, t\right) = S_{\mathrm{D}}(x) S_{\mathrm{G}}(x) \int_{s_{\mathrm{b}}}^{\infty} f(s, x, t, E) u(s, x, t) ds, \tag{71}$$

where $s_{\mathrm{b}}$ is the size at birth, $S_{\mathrm{D}}$ is the probability of seed survival during dispersal, and $S_{\mathrm{G}}$ is the probability of survival during germination.

The probability of survival during germination is assumed to depend on productivity,

$$S_{\mathrm{G}} = \frac{(P_{\mathrm{net}}/A_{\mathrm{c}})^2}{(P_{\mathrm{net}}/A_{\mathrm{c}})^2 + P_{50}^2}, \tag{72}$$

where $P_{50}$ is the productivity required to achieve 50% chance of survival.

The probability of survival during dispersal ($S_{\mathrm{D}}$) is assumed to be constant.

### 4 The Light environment

In Plant-FATE, light is the main resource for which trees compete. Taller trees shade shorter ones. Light diminishes as it passes through the canopy. The vertical light profile is affected by the height distribution of trees in the population as well as the spatial distribution of tree crowns in the canopy.

#### 4.1 Canopy layers and the PPA

We assume that position their crowns optimally in canopy gaps, in a way that minimizes overlap with other crowns. This assumption, called the perfect plasticity approximation (PPA)[15–17], leads to a spontaneous emergence of discrete canopy layers, each containing a unit crown projetcion area per unit ground area. It also leads to a convergence in crown join heights at the lower edge of each layer (Fig. 6). Real trees achieve such optimal crown placement by slightly bending their stems.

First, let us consider the first canopy layer, which closes at height $z^*$, i.e., the crown area above $z^*$ equals the ground area $A$. Therefore, $z^*$ satisfies the following identity,

$$A_{\mathrm{p},1}\left(z^*\right) + A_{\mathrm{p},2}\left(z^*\right) + \ldots = A\left(1 - f_{\mathrm{G}}\right), \tag{73}$$

where $A_{\mathrm{p},i}\left(z^*\right)$ is the crown projection area of tree $i$ at height $z^*$, $A$ is the area of the plot, $f_{\mathrm{G}}$ is the fraction of the layer that remains unoccupied due to spacing between individual crowns

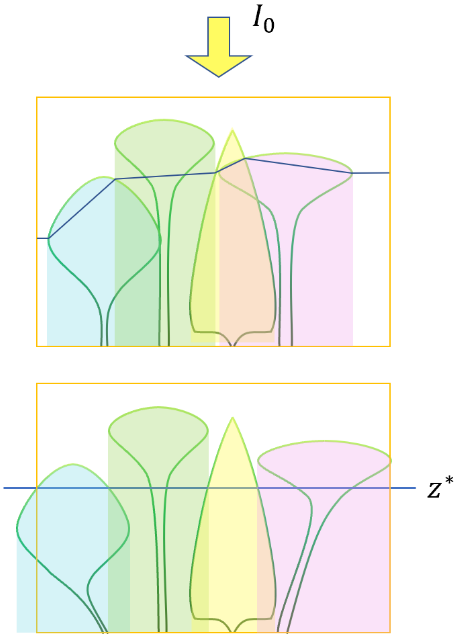

Fig. 6: **Motivating the Perfect Plasticity Approximation (PPA).** If trees can slightly bend their stems to place their crowns optimally in canopy gaps and thus avoid shading from neighbours, this would closely mirror the PPA assumption. The PPA posits that trees can strategically position their crowns in canopy gaps to ensure full light above the crown-join height z* and shading by exactly one leaf layer immediately below z*. The width of the orange box represents a unit of ground area. The transparently coloured regions indicate the shaded areas beneath each of the four trees. The crown-join height z* is calculated such that the crown area around this height equals the ground area.

(e.g., resulting from crown shyness), and the index $i$ runs over every tree in the population. For a tree of height $H$, $A_{\mathrm{p}}$ is the area enclosed by the crown's projection at height $z^*$, including the area left unfilled due to gaps. Note that the projection area reaches a maximum at $z = z_{\mathrm{m}}$, and thus $A_{\mathrm{p}}$ is given by

$$A_{\mathrm{p}}(z^*) = \begin{cases} \pi r^2(z^*), & z^* > z_{\mathrm{m}} \\ \pi r^2(z_{\mathrm{m}}), & z^* \leq z_{\mathrm{m}}, \end{cases} \tag{74}$$

with

$$\pi r^2(z) = \pi r_0^2 q(z)^2 = A_c \left(\frac{q(z)}{q_{\mathrm{m}}}\right)^2. \tag{75}$$

Writing Eq. 73 in terms of the size distribution function, we get

$$\int_{S(z^*)}^{\infty} N(s,t) A_{\mathrm{p}}(z^*, s)\, ds = A\left(1 - f_{\mathrm{G}}\right), \tag{76}$$

where $A_p(z^*, s)$ is the projected crown area at height $z^*$ of a tree of size $s$, and $N(s,t)ds$ is the total number of individuals in the plot within the size interval $[s, s+ds)$ (i.e., $N(s,t) = u(s,t)A$), and $S(z^*)$ is the size of individuals with height $z^*$. Dividing both sides by the plot area $A$, we get

$$\int_{S(z^*)}^{\infty} u(s,t) A_{\mathrm{p}}(z^*, s)\, ds = \left(1 - f_{\mathrm{G}}\right). \tag{77}$$

If there are multiple species (indexed by $k$) in the population, we must consider the contribution from all species to the crown area in the layer,

$$\sum_k \int_{S_k(z^*)}^{\infty} u_k(s,t) A_{\mathrm{p},k}(z^*, s)\, ds = \left(1 - f_{\mathrm{G}}\right).$$

The space between heights $[z^*, \infty)$ forms the first (topmost) layer of the canopy. If $z^* > 0$, the canopy is fully closed at $z = z^*$.

Now, consider the case in which the total crown area is sufficient to occupy two or more canopy layers. Let the first layer lie between $[z_1^*, \infty)$. Then, we can find a second height $z_2^*$ above which there are two fully occupied canopy layers. In general, layer $l$ has $l$ fully occupied crown layers above it. We can find $z_l^*$ by solving

$$\sum_k \int_{S_k(z_l^*)}^{\infty} u_k(s,t) A_{\mathrm{p},k}\left(z_l^*, s\right) ds = l\left(1 - f_{\mathrm{G}}\right). \tag{78}$$

Further, since trees with height lesser that $z_l^*$ (size less than $S_k(z_l^*)$) have zero projected crown area above height $z_l^*$, it follows that

$$\int_0^{S_k(z_l^*)} u_k(s,t) A_{\mathrm{p},k}\left(z_l^*, s\right) ds = \int_0^{S_k(z_l^*)} u_k(s,t) \cdot 0 \; ds = 0. \tag{79}$$

This allows us to replace the lower limit of the integral with 0,

$$\sum_k \int_0^{\infty} u_k(s,t) A_{\mathrm{p},k}\left(z_l^*, s\right) ds = l\left(1 - f_{\mathrm{G}}\right). \tag{80}$$

While Eqs. 78 and 80 give identical results, Eq. 80 is less prone to artefacts of discretization, and thus more numerically robust.

##### 4.2 Light transmission through the canopy

The number of photons absorbed per unit time by leaves in layer $l$ is given by

$$P_{\mathrm{abs},l} = I_l\left(\Delta A_{\mathrm{cp},l,1}\left(1 - e^{-kL_1}\right) + \Delta A_{\mathrm{cp},l,2}\left(1 - e^{-kL_2}\right) + \dots\right), \tag{81}$$

where $I_l$ is the light intensity (number of incident photons per unit area per unit time) at the top of layer $l$, $L_i$ is the crown LAI of tree $i$, and $\Delta A_{\mathrm{cp},l,i}$ is the filled crown projection area of tree $i$ within layer $l$. $\Delta A_{\mathrm{cp},l}$ is thus given by

$$\Delta A_{\mathrm{cp},l} = A_{\mathrm{cp}}\left(z_l^*\right) - A_{\mathrm{cp}}\left(z_{l-1}^*\right), \tag{82}$$

where $A_{\mathrm{cp}}(z)$ is the filled crown projection area of the tree at height $z$, given by

$$\begin{aligned} A_{\mathrm{cp}}(z) &= \begin{cases} \pi r^2(z)(1 - f_{\mathrm{g}}), & z > z_{\mathrm{m}} \\ \pi r^2\left(z_{\mathrm{m}}\right)\left(1 - f_{\mathrm{g}}\right) + \left(\pi r^2\left(z_{\mathrm{m}}\right) - \pi r^2(z)\right) f_{\mathrm{g}}, & z \leq z_{\mathrm{m}} \end{cases} \\ &= \begin{cases} A_{\mathrm{c}}\left(\frac{q(z)}{q_{\mathrm{m}}}\right)^2 (1 - f_{\mathrm{g}}), & z > z_{\mathrm{m}} \\ A_{\mathrm{c}} - A_{\mathrm{c}}\left(\frac{q(z)}{q_{\mathrm{m}}}\right)^2 f_{\mathrm{g}}, & z \leq z_{\mathrm{m}}. \end{cases} \end{aligned} \tag{83}$$

For the first layer with $l = 1$, we define $A_{\text{cp}}(z_0^*) = A_{\text{cp}}(\infty) = 0$.

Writing Eq. 81 in terms of the size distribution, we get

$$P_{\text{abs},l} = I_l \sum_k \int_0^\infty N_k(s,t) \Delta A_{\text{cp},l,k}(s) \left(1 - e^{-kL_k(s)}\right) ds, \tag{84}$$

The average absorbed light intensity is then

$$I_{\text{abs},l} = \frac{P_{\text{abs},l}}{A} = I_l \sum_k \int_0^\infty u_k(s,t) \Delta A_{\text{cp},l,k}(s) \left(1 - e^{-kL_k(s)}\right) ds. \tag{85}$$

The layer-average fraction of absorbed light is then

$$f_{\text{abs},l} = \frac{I_{\text{abs},l}}{I_l} = \sum_k \int_0^\infty u_k(s,t) \Delta A_{\text{cp},l,k}(s) \left(1 - e^{-kL_k(s)}\right) ds \tag{86}$$

The average light intensity at the top of layer $l + 1$ is the light intensity transmitted through layer $l$, given by

$$I_{l+1} = I_l \left(1 - f_{\text{abs},l}\right) \tag{87}$$

Dividing both sides by the incident radiation at the top of the canopy $I_0$, and defining the canopy openness of layer $l$ as $C_l = I_l / I_0$, we get a recursion for calculating the vertical light profile,

$$\begin{aligned} C_1 &= 1, \\ C_{l+1} &= C_l \left(1 - f_{\text{abs},l}\right). \end{aligned} \tag{88}$$

In summary, the light environment is specified by the the canopy layer boundaries $\{z_1^*, z_2^*, ...\}$ and the fraction of top-canopy light intensity available in each layer $\{1, C_1, C_2, ...\}$.

#### 4.3 Light availability for photosynthesis

For calculating whole-plant photosynthesis, there are two possibilities: (i) calculate asismilation rates separately for each layer and sum them up over layers, or (ii) calculate average light intensity across layers and use it to calculate an averge assimilation rate. Since layers can change continuously, causing leaves to move from one layer to another, the first option requires keeping track of acclimated photosynthetic capacity at each height $z$. Since this is computationally extremely inefficient, we take the second option, which is computationally efficient at the expense of only a negligible difference in the result.

Thus, we calculate the weighted average of the light intensity as follows,

$$I_{\text{avg}} = \sum_l \frac{\Delta A_{\text{cp},l}}{A_{\text{c}}} I_l. \tag{89}$$

where the layer weight is the the proportion of total crown projection area in the layer.

The fraction of absorbed light $f_{\text{apar}}$ is calculated frm the crown LAI,

$$f_{\text{apar}} = 1 - e^{-kL}, \tag{90}$$

where $k$ is the light extinction coefficient.

### A Appendix A: Justification of constant Huber Value within the crown

The drop in water potential along the stem and branches of the plant due to transpiration can be expressed by the differential equation (see SI in P-hydro paper)

$$\frac{d\psi}{dh} = -\frac{\eta E}{\kappa_{\mathrm{s}}(\psi, h)\nu_{\mathrm{H}}} - \rho g. \tag{91}$$

We assume that the maximum conductivity $\kappa(h)$ of xylem segments increases depends on $h$ due to xylem tapering. This maximum conductivity is reduced due to embolism such that the realised conductivity is $\kappa_{\mathrm{s}}(\psi, h) = \kappa(h)P(\psi)$, where $P(\psi)$ is the vulnerability curve. We assume that $\psi_{50}$, which is a parameter in the vulnerability curve, does not depend on $h$.

Furthermore, we assume that the xylem gets wider as we move from the leaf petiole towards the stem base. Following a continuous analogue of the WBE model, we assume that

$$\kappa(h) = \kappa_{\mathrm{p}} e^{\sigma_{\mathrm{p}}(l-h)}, \tag{92}$$

where $l$ is the distance from the ground to the petiole. Substituting in Eq. 91, we get

$$\begin{aligned} \int_{\psi_s}^{\psi_l} P(\psi) d\psi &= -\int_0^l \frac{\eta E}{\kappa_{\mathrm{p}}\nu_{\mathrm{H}} e^{\sigma_{\mathrm{p}}(l-h)}} dh - \rho_{\mathrm{w}} g l_{\perp} \\ &= -\frac{\eta E}{\kappa_{\mathrm{p}}\sigma_{\mathrm{p}}\nu_{\mathrm{H}}}\left(1 - e^{-\sigma_{\mathrm{p}}\left(z + r\tan\left(\frac{\theta}{2}\right)\right)}\right) - \rho_{\mathrm{w}} g z. \end{aligned} \tag{93}$$

Under normal functioning, there is minimal embolism, so $P(\psi) \approx 1$ and thus integral on the LHS is $\psi_{\mathrm{l}} - \psi_{\mathrm{s}} = -\Delta\psi$. We further ignoring gravitational potential drop. Consider three possibilities about how $\nu_{\mathrm{H}}$ and $\Delta\psi$ vary within a plant.

1. The Huber Value $\nu_{\mathrm{H}}$ remains constant across the plant, and $\Delta\psi$ varies as

$$\Delta\psi(z) = \Delta\psi_{\infty}\left(1 - e^{-\sigma_{\mathrm{p}}\left(z + r\tan\left(\frac{\theta}{2}\right)\right)}\right), \tag{94}$$

with

$$\Delta\psi_{\infty} = \frac{\eta E}{\kappa_{\mathrm{p}}\sigma_{\mathrm{p}}\nu_{\mathrm{H}}}. \tag{95}$$

2. The potential drop $\Delta\psi$ remains constant throughout the plant, which necessitates a change

in Huber Value as

$$\nu_{\mathrm{H}} = \nu_{\mathrm{H},\infty}\left(1 - e^{-\sigma_{\mathrm{p}}\left(z + r\tan\left(\frac{\theta}{2}\right)\right)}\right), \tag{96}$$

with

$$\nu_{\mathrm{H},\infty} = \frac{\eta E}{\kappa_{\mathrm{p}}\sigma_{\mathrm{p}}\Delta\psi}. \tag{97}$$

3. A combination of the above two, where neither is constant. There is evidence that the leaf water potential does change with height within a plant, and there is no adequate data to comment on Huber Value. We therefore assume the first possibility, as it leads to the simplest allometric scaling equations.

### B Supplementary figures

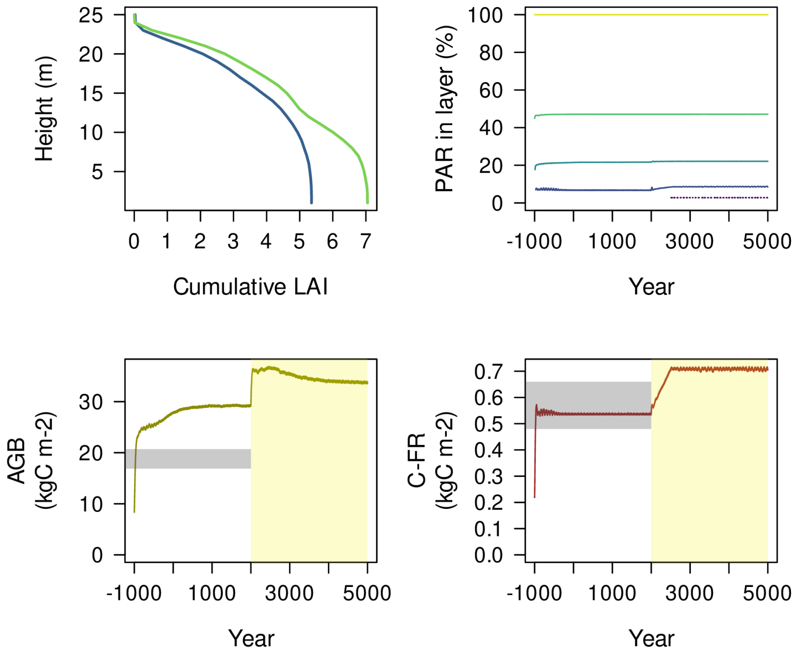

Fig. S1: **Additional emergent community properties predicted by our model under elevated and ambient CO2 conditions.**

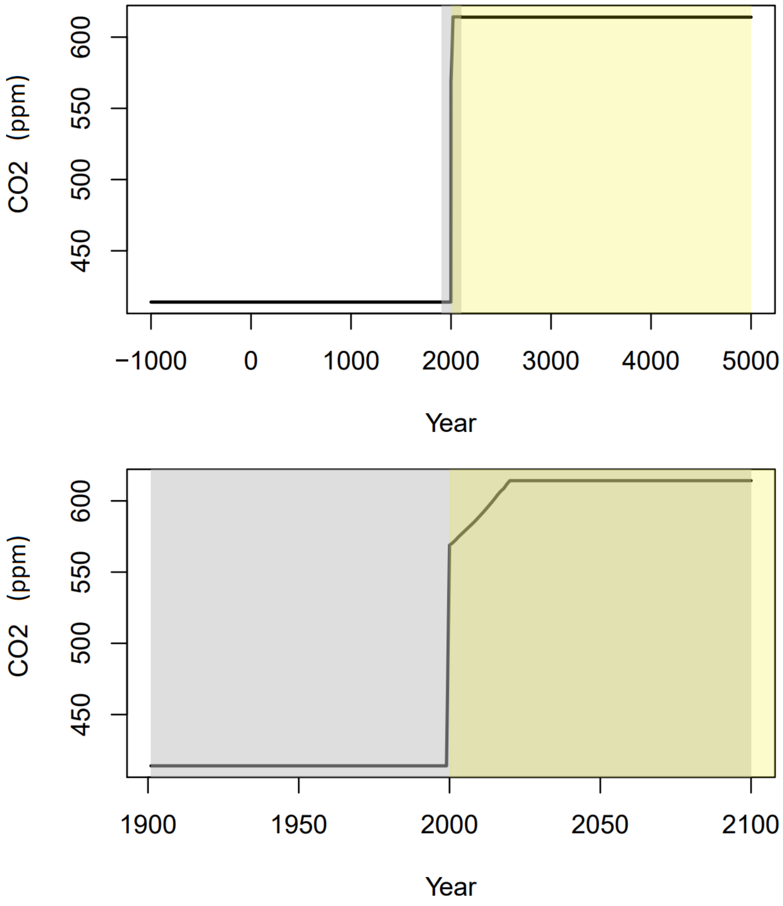

Fig. S2: **Time series of atmospheric CO2 forcing used for our model runs.**

10. I. A. Janssens, H. Lankreijer, G. Matteucci, A. S. Kowalski, N. Buchmann, D. Epron, K. Pilegaard, W. Kutsch, B. Longdoz, T. Grünwald, L. Montagnani, S. Dore, C. Rebmann, E. J. Moors, A. Grelle, Ü. Rannik, K. Morgenstern, S. Oltchev, R. Clement, J. Guðmundsson, S. Minerbi, P. Berbigier, A. Ibrom, J. Moncrieff, M. Aubinet, C. Bernhofer, N. O. Jensen, T. Vesala, A. Granier, E. D. Schulze, A. Lindroth, A. J. Dolman, P. G. Jarvis, R. Ceulemans, and R. Valentini. Productivity overshadows temperature in determining soil and ecosystem respiration across European forests: RESPIRATION ACROSS EURO-

PEAN FORESTS. *Global Change Biology*, 7(3):269–278, March 2001. ISSN 1354-1013. doi: 10.1046/j.1365-2486.2001.00412.x.

17. E. S. Weng, S. Malyshev, J. W. Lichstein, C. E. Farrior, R. Dybzinski, T. Zhang, E. Shevliakova, and S. W. Pacala. Scaling from individuals to ecosystems in an Earth System Model using a mathematically tractable model of height-structured competition for light. preprint, December 2014.
